# Dynamic phosphatase-recruitment controls B-cell selection and oncogenic signaling

**DOI:** 10.1101/2023.03.13.532151

**Authors:** Jaewoong Lee, Mark E. Robinson, Ruifeng Sun, Kohei Kume, Ning Ma, Kadriye Nehir Cosgun, Lai N. Chan, Etienne Leveille, Huimin Geng, Vivasvan S. Vykunta, Brian R. Shy, Alexander Marson, Samuel Katz, Jianjun Chen, Elisabeth Paietta, Eric Meffre, Nagarajan Vaidehi, Markus Müschen

## Abstract

Initiation of B-cell receptor (BCR)^1^ signaling, and subsequent antigen-encounter in germinal centers^2,3^ represent milestones of B-lymphocyte development that are both marked by sharp increases of CD25 surface-expression. Oncogenic signaling in B-cell leukemia (B-ALL)^4^ and lymphoma^5^ also induced CD25-surface expression. While CD25 is known as an IL2-receptor chain on T- and NK-cells^6–9^, the significance of its expression on B-cells was unclear. Our experiments based on genetic mouse models and engineered patient-derived xenografts revealed that, rather than functioning as an IL2-receptor chain, CD25 expressed on B-cells assembled an inhibitory complex including PKCδ and SHIP1 and SHP1 phosphatases for feedback control of BCR-signaling or its oncogenic mimics.

Recapitulating phenotypes of genetic ablation of PKCδ^10^–^12^, SHIP1^13,14^ and SHP1^14, 15,16^, conditional CD25-deletion decimated early B-cell subsets but expanded mature B-cell populations and induced autoimmunity. In B-cell malignancies arising from early (B-ALL) and late (lymphoma) stages of B-cell development, CD25-loss induced cell death in the former and accelerated proliferation in the latter. Clinical outcome annotations mirrored opposite effects of CD25-deletion: high CD25 expression levels predicted poor clinical outcomes for patients with B-ALL, in contrast to favorable outcomes for lymphoma-patients. Biochemical and interactome studies revealed a critical role of CD25 in BCR-feedback regulation: BCR-signaling induced PKCδ-mediated phosphorylation of CD25 on its cytoplasmic tail (S^268^). Genetic rescue experiments identified CD25-S^268^ tail-phosphorylation as central structural requirement to recruit SHIP1 and SHP1 phosphatases to curb BCR-signaling. A single point mutation CD25^S268A^ abolished recruitment and activation of SHIP1 and SHP1 to limit duration and strength of BCR-signaling. Loss of phosphatase-function, autonomous BCR-signaling and Ca^2+^-oscillations induced anergy and negative selection during early B-cell development, as opposed to excessive proliferation and autoantibody production in mature B-cells. These findings highlight the previously unrecognized role of CD25 in assembling inhibitory phosphatases to control oncogenic signaling in B-cell malignancies and negative selection to prevent autoimmune disease.

---

Immunoreceptor tyrosine-based activation (ITAM)^17^ or inhibition (ITIM)^18^ motifs function as conserved sequences in the signaling chains of B-cells receptors (BCR) to activate tyrosine kinases to amplify or phosphatases to terminate BCR signaling, respectively. However, phosphorylated ITAM and ITIM motifs can attract and activate tyrosine kinases and phosphatases only within a range of ∼5 nm^19^. Hence, additional recruitment mechanisms for long-range recruitment of cytoplasmic kinases and phosphatases are necessary to concentrate their local abundance in proximity of BCR molecules. Central molecules that amplify BCR-signaling, including BTK and AKT kinases, VAV1 and SOS1 guanine nucleotide exchange factors, as well as PLCγ1 and PLCγ2, are recruited based on interactions between their PH-domains and PIP3 in lipid rafts^20,21^. In addition, the conserved intracellular loop of Ifitm3 serves as an alternative structure to recruit activating kinases to BCR-signaling complexes within lipid rafts^22^. While mechanisms of local recruitment of activating kinases in lipid rafts have been extensively studied, how inhibitory phosphatases are recruited to curtail local signaling activity remained elusive. Here we identified CD25-S^268^ tail-phosphorylation as the central structural element of phosphatase recruitment and feedback control of BCR-signaling in activated and transformed B-cells.

This was unexpected because CD25 was previously known as one of the three chains of the IL2-receptor on T-cells and NK-cells^6–9^. While the IL2Rβ and the common γ-chain (IL2Rγ) are essential for cytoplasmic signal transduction^23,24^, CD25 does not contribute to IL2-signal transduction. Instead, CD25 interacts with IL2 and induces a small conformational change of IL2, which increases binding affinity to the IL2Rβ-chain^8,9^. Loss of IL2Rβ and IL2Rγ results in profound defects of T-cell and NK-cell production. In contrast to IL2Rβ and IL2Rγ, CD25-deficiency causes lymphoproliferation and autoimmunity^25–27^. Instead of defective IL2-signaling, the phenotype of CD25-deficient humans and mice resembled B-cell hyperactivation and autoimmunity in mice lacking functional PKCδ^10–12^ or inhibitory SHIP1^13,14^ and SHP1^14,15,16^-phosphatases. Interestingly, polymorphisms of the *IL2RA* gene encoding CD25 were associated with increased risk of multiple autoimmune diseases (type 1 diabetes^28^, multiple sclerosis^29,30^, juvenile idiopathic arthritis^31^, systemic lupus erythematosus^32^) and predisposition to B-cell acute lymphoblastic leukemia (B-ALL)^33^.

All three chains of the IL2 receptor are endocytosed together after IL2 binding: IL2Rβ and IL2Rγ chains are targeted to late endosomes for degradation, while CD25 chains are segregated to early endosomes for recycling to the cell membrane^34,35^. Consistent with different fates in IL2-activated T cells, the half-life of CD25 chain (>40 hours) is much longer than IL2Rβ and IL2Rγ (55-70 minutes) chains^34,35^. Its lack of contribution to IL2 signal transduction, asynchronous half-life and divergent intracellular trafficking suggest that CD25 has functions in lymphocyte signaling that are independent from the other two IL2-receptor chains.

While B-cells develop in normal numbers in the absence of IL2-signaling^36^, CD25 surface levels are strongly upregulated at two central milestones during B-cell development, namely the first assembly of a functional pre-BCR signaling complex^1^ and subsequent antigen-encounter of the BCR expressed on mature B-cells^2,3^. In addition, CD25 is expressed at high levels in multiple B-cell malignancies^4,5^, the significance of which was not clear. Here we identified CD25 as a negative feedback regulator of (pre-)BCR-signaling during normal B-cell development, as well as of oncogenes in B-cell malignancies that mimic BCR-downstream signaling pathways. While the β- and γ-chains of the IL2-receptor can be repurposed to become part of other cytokine receptors^37,38^, CD25 represents the first example of a cytokine receptor chain that can be recycled to recruit an inhibitory complex for feedback control of BCR-signaling or its oncogenic mimics.

## CD25 surface expression marks milestones of B-cell development and oncogenic signaling

CD25 surface levels are markedly upregulated at two central milestones during B-cell development, namely the first assembly of a functional pre-BCR signaling complex^1^ and subsequent antigen-encounter of the BCR expressed on mature B-cells^2,3^. To determine how CD25 is mechanistically linked to the initiation of pre-BCR signaling during early and antigen-encounter during late B-cell development, we performed genetic experiments to model these two milestones. *Rag2*^-/-^ pro-B cells lack the ability to rearrange V_H_, D and J_H_ segments for expression of a functional Ig μ-heavy chain (μHC) as part of the pre-BCR^39^. *Blnk*-deficient pre-B cells express a functional μHC but lack pre-BCR signal transduction^40^. In these models, reconstitution of μHC-expression and pre-BCR signal transduction induced marked CD25 surface expression (**Extended Data Figure 1a-c**). Likewise, BCR-engagement, to model antigen encounter in mature B-cells, induced CD25 surface expression, which was sensitive to treatment with ibrutinib, a small molecule inhibitor of the proximal BCR-downstream tyrosine kinase BTK (**Extended Data Figure 1d**). Consistent with antigen-driven upregulation of CD25 surface expression during late stages of B-cell development, antigen-experienced human tonsillar germinal center and memory B-cells expressed CD25 at higher levels than naïve B-cells (**Extended Data Figure 1e**). Since some subtypes of B-cell leukemia (B-ALL)^4^ and lymphoma^5^ express high levels of CD25 on the surface, we tested whether oncogenic signaling in B-ALL and B-cell lymphoma promotes CD25 expression. The BCR-ABL1 kinase functions as an oncogenic mimic of pre-BCR signaling in B-ALL^41^ and promoted BTK-dependent CD25-upregulation on transformed pre-B cells (**Extended Data Figure 1f**). Likewise, CARD11^L225LI^, IKK2^S177E^ and MYD88^L265P^ oncogenes promote oncogenic NF-κB signaling^42–44^ and upregulated CD25 when expressed in tonsillar memory B-cells (**Extended Data Figure 1g**). Consistent with strong induction of mRNA and surface expression of CD25 by IKK2^S177E^ and MYD88^L265P^, multiple NF-κB subunits bound to the *CD25* promoter (**Extended Data Figure 1j-k**). In patient-derived B-ALL and lymphoma samples, oncogenic signaling induced CD25 surface expression only in a fraction of malignant cells (**Extended Data Figure 1f-i**). Comparing patient-derived CD25^+^ and CD25^-^ B-ALL cases in a clinical trial (ECOG E2993), CD25-expression was strongly associated with an NF-κB gene expression program (**Extended Data Figure 1l-m**). Comparing flow-sorted CD25^High^ and CD25^Low^ B-ALL cells from three patients, confirmed the association between CD25-expression and an NF-κB transcriptional program. In addition, CD25^High^ B-ALL cells were small and quiescent despite their increased STAT5, AKT and ERK signaling activity (**Extended Data Figure 2a-e**). Reduced proliferation and colony formation activity of CD25^High^ B-ALL cells was paralleled by resistance to chemotherapy (vincristine; **Extended Data Figure 2f-g**). Likewise, leukemia-initiation from CD25^High^ cells was delayed compared to CD25^Low^ B-ALL cells, however, both populations achieved complete penetrance of the disease in transplant recipients (**Extended Data Figure 2h**). These results highlight that CD25 not only marks critical milestones during normal B-cell development but also drug-resistant and quiescent subpopulations in patient-derived B-ALL.

## Opposite effects of CD25-deletion during early and late stages of B-cell development

To examine a potential mechanistic role of CD25 during B-cell development and in B-cell malignancies, we developed a conditional mouse model for B-cell-specific (*Cd25*^fl/fl^ x *Mb1*-Cre) and tamoxifen-inducible (*Cd25*^fl/fl^ x *Mb1*-Cre^ERT2^) deletion of CD25. Strikingly, B-cell-specific ablation of CD25 profoundly reduced the pool of bone marrow pre-B and immature B-cells, while mature splenic B-cells and peritoneal cavity B1-cell populations were substantially expanded (**Figure 1a-c, Extended Data Figure 3**). Inducible deletion of CD25 *in vitro* caused rapid depletion and cell death in cultured bone marrow pre-B cells as opposed to expansion of splenic mature B-cell cultures (**Figure 1d-e**). Likewise, pre-B cell-derived B-ALL cells (transformed by *NRAS*^G12D^ and *BCR-ABL1* oncogenes) underwent cell cycle arrest, lost colony formation ability, and died under cell culture conditions upon deletion of CD25. In contrast, loss of CD25 accelerated proliferation and colony formation of mature B-cell lymphoma cells (*Card11*^L232LI^; **Figure 1f-i**).

**Figure 1:**
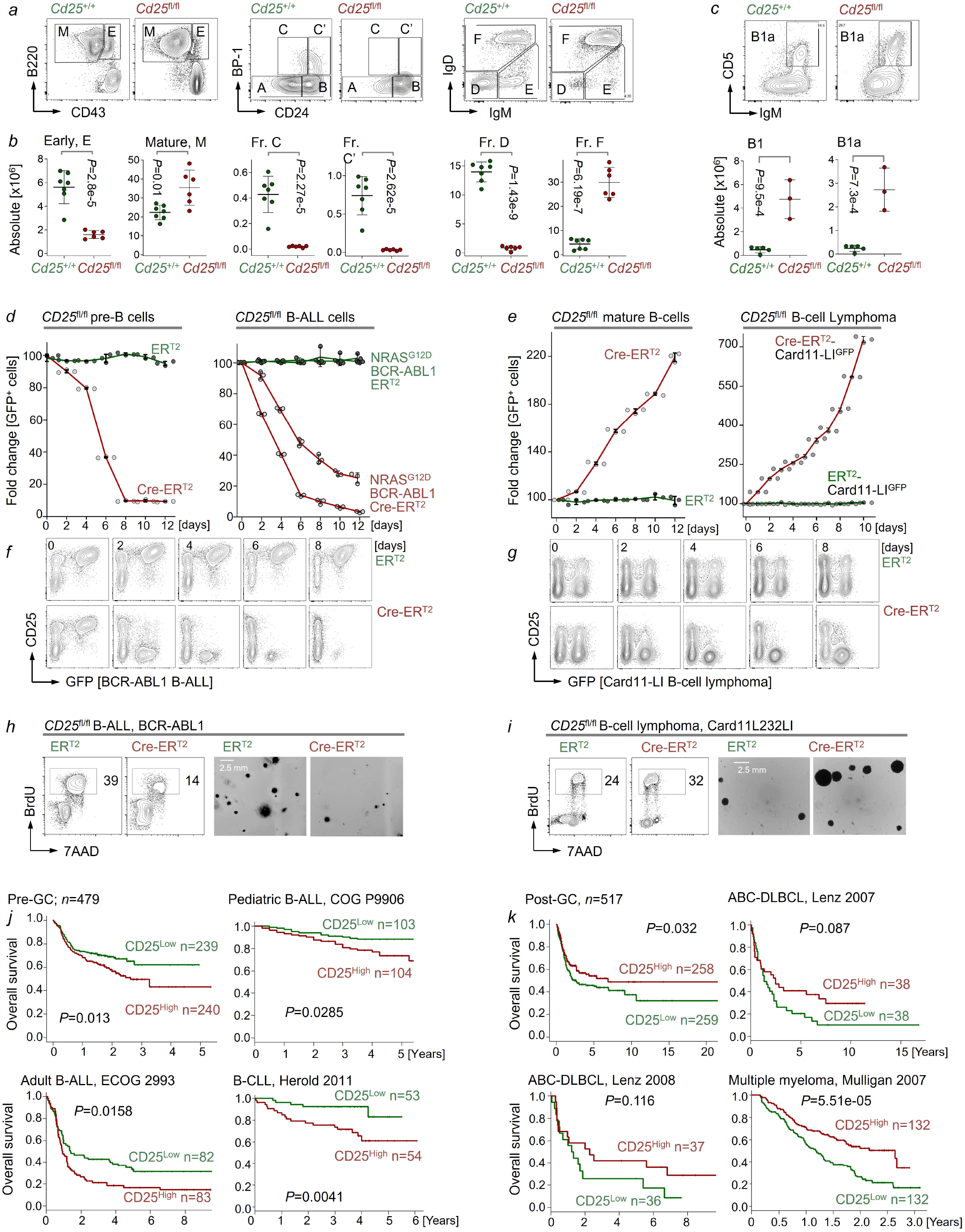
CD25 is essential during early stages of B-cell development but negatively regulates mature B-cell proliferation. **a-b**, Hardy fractions of B cell subsets isolated from bone marrow of *Cd25*^+/+^ *Mb1*-Cre (n=7) and *Cd25*^fl/fl^ *Mb1*-Cre (n=6) littermates were analyzed by flow cytometry; representative plots are shown (**a**). Among Nk1.1^-^ and Gr1^-^ populations, absolute cell counts of early progenitor (“E”, B220^+^, CD43^+^), including fraction C (Fr. C, B220^+^, CD43^+^, CD24^low^, BP-1^high^), fraction C’ (Fr. C’, B220^+^, CD43^+^, CD24^high^, BP-1^high^), fraction D (Fr. D, B220^+^, CD43^-^, IgM^low^, IgD^low^) and mature B-cells (“M”, B220^+^, CD43^-^), including fraction F (Fr. F, B220^+^, CD43^-^, IgM^high^, IgD^high^) cells were plotted (**b**). Absolute cell counts of B1 (CD19^+^, Mac1^+^, IgM^+^) and B1a (CD19^+^, CD5^+^, IgM^+^) cells in the peritoneal cavity from *Cd25*^+/+^ *Mb1*-Cre (n=5) and *Cd25*^fl/fl^ *Mb1*-Cre (n=3) littermates were plotted with representative flow cytometric analysis (**c**). IL7-dependent *Cd25*^fl/fl^ pre-B cells or B-ALL cells retrovirally transformed with either NRAS^G12D^ or BCR-ABL1 were transduced with 4-hydroxytamoxifen (4-OHT)-inducible Cre-ER^T2^-GFP or ER^T2^-GFP empty vector control. Percentages of GFP^+^ cells were measured by flow cytometry (n=3) at different time points following 4-OHT treatment (**d**). *Cd25*^fl/fl^ mature B-cells purified from splenocytes by negative selection were retrovirally transformed with Card11^L232LI^ to develop mature B-cell lymphoma and then transduced with 4-OHT-inducible Cre-ER^T2^-GFP or ER^T2^-GFP in the presence of IL4 (10 ng/ml), BAFF (100 ng/ml) and CD40L (1 μg/ml). In *Cd25*^fl/fl^ splenic mature B-cells or lymphoma cells, percentages of GFP^+^ cells were measured by flow cytometry (n=3) at different time points following 4-OHT treatment (**e**). The effect of inducible deletion of CD25 in *Cd25*^fl/fl^ B-ALL (**f**) and lymphoma cells (**g**) was monitored by flow cytometry at different time points after 4-OHT treatment. B-ALL (BCR-ABL1) cells derived from *Cd25*^fl/fl^ pre-B cells (**h**), and B-cell lymphoma (Card11^L232LI^) derived from *Cd25*^fl/fl^ mature splenic B-cells (**i**) were transduced with 4-OHT-inducible Cre-ER^T2^ or ER^T2^ empty vector control. Cell cycle analyses were performed by measuring BrdU incorporation in combination with 7AAD staining following 4-OHT treatment for 2 days, and the percentages of cells in S phase are indicated (**h-i**, *left*). Cells were plated in semi-solid methylcellulose in the presence of 4-OHT for colony-forming assay for 7 days and colony-forming capacity was assessed (**h-i**, *right*). Patients with pediatric B-ALL (n=207, P9906), adult B-ALL (n=165, E2993), B-CLL (n=107, GSE22762) as “pre-GC” (top-left, total n=479; **j**) and ABC-DLBCL (n=76, Lenz 2007; n=73, Lenz 2008), Multiple myeloma (n=264, Mulligan 2007) as “post-GC” (top-left, total n=517; **k**) were divided into two groups based on higher or lower than median levels of *CD25* mRNA. Overall survival was estimated in the two groups by Kaplan-Meier survival analyses. Log-rank test was used to assess statistical significance. In **b, c**, Statistical significance was assessed by two-tailed *t*-test (means ± s.d.). In **f, g, h, i**, Representative plots and images are shown (n=3). In **h, i**, Scale bar, 2.5 mm.

Clinical outcome annotations for CD25 expression levels in clinical cohorts mirrored these differences between B-cell malignancies derived from early and late stages of B-cell development. Studying gene expression data from six clinical trials for patients with B-cell malignancies derived from early or pre-GC (B-ALL, CLL) and late or post-GC (DLBCL, multiple myeloma) stages of B-cell development, we found that high CD25 expression levels predicted poor clinical outcomes for patients with B-ALL and CLL, in contrast to favorable outcomes for patients with DLBCL and myeloma (**Figure 1j-k, Extended Data Figure 2a-b**). These results suggest that CD25 has opposite functions in regulating survival and proliferation during early and late B-cell development and that these differences also manifest in B-cell tumors of distinct cellular origins.

## CD25-deficient mature B-cells are hyperactive and give rise to autoimmunity

Functional studies of the enlarged mature B-cell compartment revealed that deletion of CD25 during early B-cell development (*Mb1*-Cre) and in germinal centers (*Aicda*-Cre) resulted in formation of spontaneous germinal centers (**Figure 2a-c**) with increased immunoglobulin serum levels (**Figure 2d**). When transplanted into NSG mice lacking endogenous B-cells, CD25-deficient mature B-cells expanded at a 9-fold increased rate compared to mature B-cells from wildtype littermates (**Figure 2e**). Hyperproliferation of CD25-deficient mature B-cells in transplant recipient mice was paralleled by production of antinuclear autoantibodies (**Figure 2f**). These lupus-like features of B-cell-specific CD25-deletion phenocopied outcomes of germinal center-specific (*Aicda*-Cre) deletion of the inhibitory SHIP1-phosphatase^45^. A closer examination of immunization experiments revealed that, NP-specific IgM serum levels were substantially increased upon B-cell-specific ablation of CD25, whereas NP-specific serum IgG levels were decreased compared to mice with CD25^+/+^ B-cells (**Figure 2d**). An *in vitro* class-switch recombination (CSR) assay revealed that concurrent increases of IgM with reduced IgG-production from CD25-deficient B-cells was the result of defective class-switching. Since class-switch recombination is negatively regulated by PI3K-mediated FOXO1-inactivation^46^, we repeated the CSR-assay with two PI3K small molecule inhibitors (**Figure 2e**). Interestingly, PI3K-inhibitors rescued defective CSR, suggesting that loss of CD25 results in a net increase of PI3K-activity in CD25-deficient B-cells. These results are consistent with an essential role of CD25 in enabling phosphatidylinositol-phosphatase SHIP1 activity, a central negative regulator of PI3K-signaling^47^.

**Figure 2:**
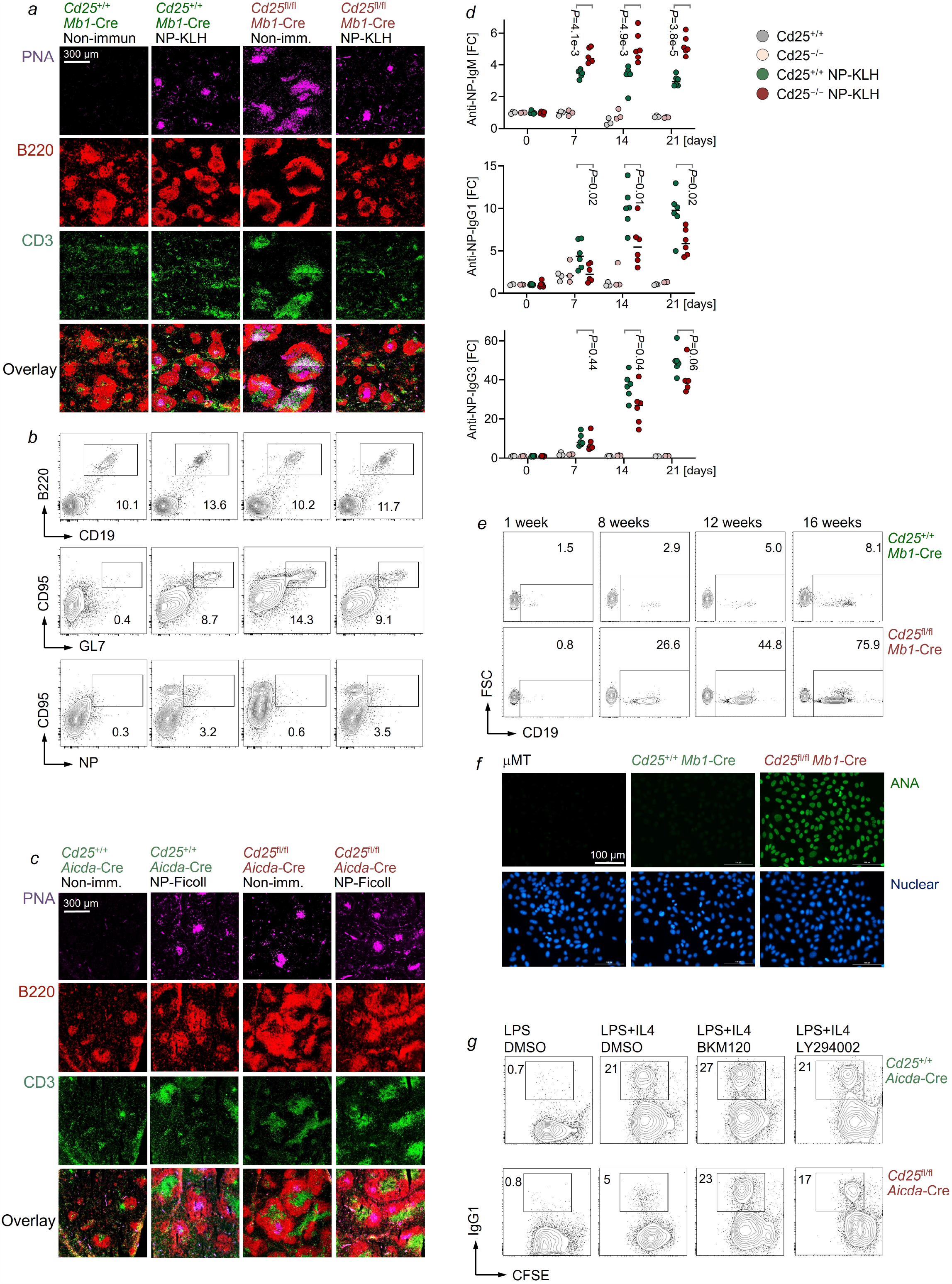
CD25 prevents formation of spontaneous germinal centers and expansion of autoreactive B-cells. *Cd25*^+/+^ *Mb1*-Cre and *Cd25*^fl/fl^ *Mb1*-Cre mice were immunized intraperitoneally with 0.5 mg of NP-KLH (n=6 per group) or vehicle (*Cd25*^+/+^, n=6; *Cd25*^fl/fl^, n=3) on day 0 and day 7 (**a-b**). Spleens were collected on day 12 after initial immunization and subjected to immunofluorescence staining of tissue sections with B220, CD3 and peanut agglutinin (PNA). Scale bar, 300 µm (**a**). Splenocytes harvested from *Cd25*^+/+^ *Mb1*-Cre and *Cd25*^fl/fl^ *Mb1*-Cre mice were analyzed by flow cytometry 12 days post-immunization for germinal center (GC)-associated markers CD95, GL7 and NP among the B220^+^ CD19^+^ population to identify GC-B cells (**b**). Representative flow cytometry plots are shown with percentages of populations. *Cd25*^+/+^ *Aicda*-Cre and *Cd25*^fl/fl^ *Aicda*-Cre mice were immunized intraperitoneally with 0.1 mg of NP-Ficoll (n=6 per group) or vehicle (n=3 per group) on day 0 (**c-d**). Spleens were collected on day 21 after initial immunization and subjected to immunofluorescence staining of tissue sections with B220, CD3 and PNA. Scale bar, 300 µm (**c**). Titers of anti-NP IgM, IgG_1_ and IgG_3_ in sera from *Cd25*^+/+^ *Aicda*-Cre and *Cd25*^fl/fl^ *Aicda*-Cre mice were assessed by ELISA at serial time points post-immunization (**d**). 10 million splenic mature B-cells from *Cd25*^+/+^ *Mb1*-Cre or *Cd25*^fl/fl^ *Mb1*-Cre were injected into NSG mice (n=7 per group). Transplanted B cells in NSG mice were monitored by flow cytometry in peripheral blood samples that were taken 1, 8, 12 and 16 weeks after injection and representative plots are shown (**e**). Sera derived from 3-month-old *Cd25*^+/+^ *Mb1*-Cre, *Cd25*^fl/fl^ *Mb1*-Cre, or µMT mice that lack endogenous mature B-cells, were used at a dilution of 1:200 for the immunofluorescent analysis of antinuclear antibodies (ANA; n=4). Scale bar, 100 µm (**f**). B-cells enriched by negative selection from *Cd25*^+/+^ *Aicda*-Cre and *Cd25*^fl/fl^ *Aicda*-Cre mice were studied in class-switching assays and stained with CFSE (10 µM) after stimulation with LPS (25 µg/ml), IL4 (10 ng/ml), or CD40L (1 µg/ml) and 4-OHT treatment for 3 days. IgG_1_ expression and dilution of CFSE dye was assessed by flow cytometry in the presence of the PI3K-inhibitors BKM120 (1 µM) or LY294002 (5 µM). Percentages of IgG1^+^ cells are indicated in each plot. Data are representative of three independent experiments. In **a, c, g**, Representative images are shown.

## CD25 is essential for activation of inhibitory phosphatases

Since B-cell-specific deletion of CD25 recapitulated multiple features of SHIP1-loss, including defective negative regulation of PI3K-activity and CSR, we hypothesized that CD25 serves a critical role in inhibitory signaling of SHIP1 and potentially other phosphatases. To elucidate a potential role of CD25 in inhibitory signaling, we performed global phosphoproteomic and by mass spectrometry analyses to identify changes of tyrosine and serine/threonine phosphorylation events upon acute deletion of CD25. Interestingly, the most prominent phosphorylation increases were observed among activating molecules in the proximal BCR-signal transduction chain, including the ITAM-motif of CD79A, SYK, BLNK, VAV1, PLCγ2, FYN, SASH3, CIN85-, PKCδ and NF-κB-activating molecules (MYD88, TRAF2, TRAF4, TRAF6, PKCβ, MALT1). Strikingly, phosphorylation events that are indicative of phosphatase-activation were most prominently lost in response to CD25-deletion. These negative changes included the ITIM-motifs of the inhibitory receptors CD22 and BTLA, phosphorylation events that reflect activation of SHIP1, SHP1, SHIP2 phosphatases, the PKCδ-adapter RACK1, the inhibitory SHIP1 adapters DOK1, DOK3, as well as SH3BP5, the inhibitor of the BCR-kinase BTK (**Figure 3a-b**). Validation experiments by co-IP corroborated that CD25 may directly interact or be part of a larger complex with the BCR signaling chain CD79B, the inhibitory phosphatases SHIP1 and SHP1 as well as PKCδ and its adapter RACK1 (**Figure 3c**). These results in murine B-ALL cells suggest that acute Cre-mediated deletion of CD25 causes imbalances between massive increases of activators of BCR-downstream signaling and near-complete loss of inhibitory phosphatase signaling.

**Figure 3:**
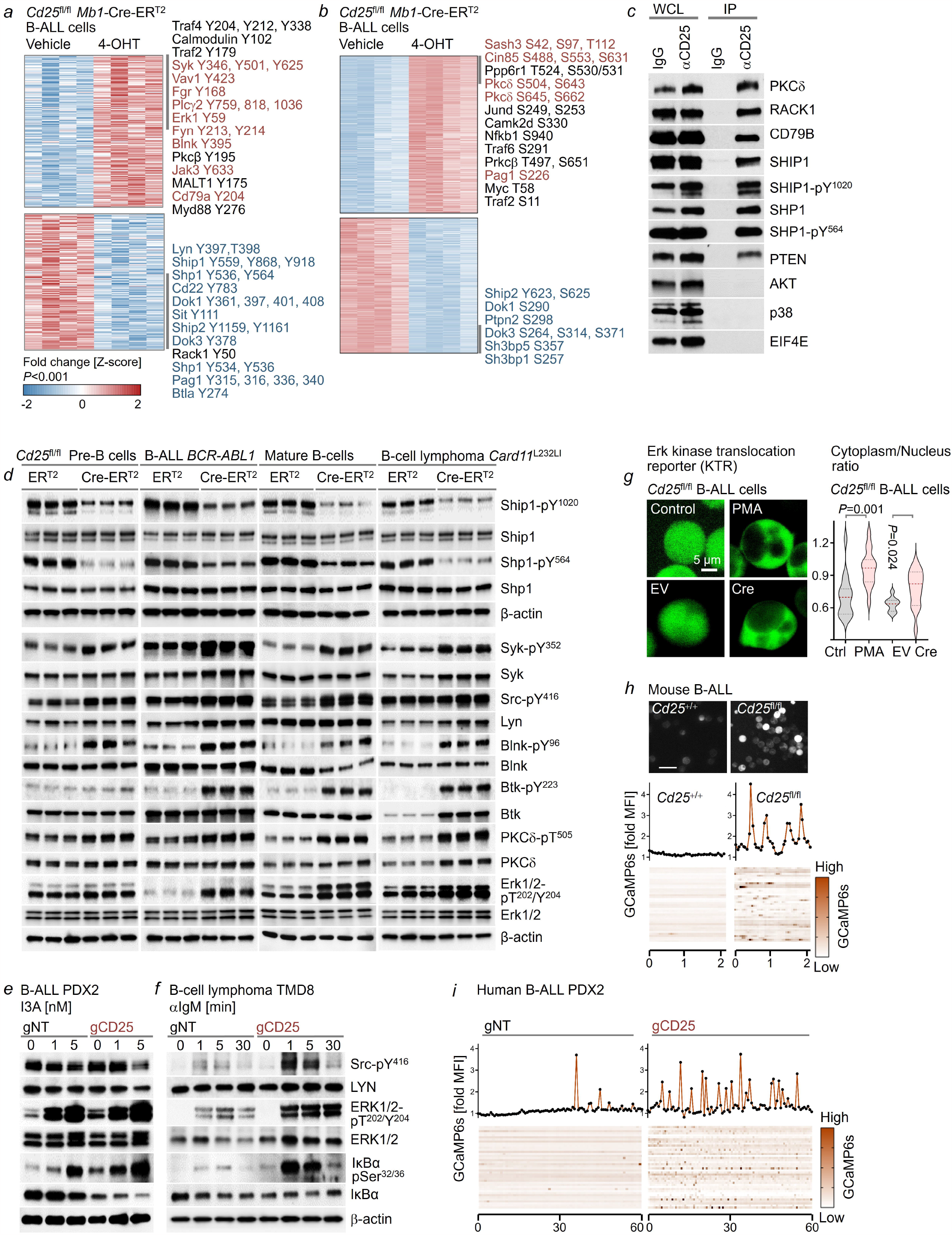
CD25 is required for activation of inhibitory phosphatases to curb signaling strength of the BCR or its oncogenic mimics. *Cd25*^fl/fl^ *Mb1*-Cre-ER^T2^ B-ALL cells transformed with BCR-ABL1 were treated with 4-OHT or vehicle for 2 days (n=4). Levels of phosphorylated proteins, determined by mass spectrometry, with phospho-tyrosine residues (**a**) or phospho-serine/threonine residues (**b**) were ranked by log_2_-transformed fold change (moderated *t*-statistic). Co-immunoprecipitation was performed in *BCR-ABL1-*transformed B-ALL cells, using anti-CD25 antibodies covalently cross-linked with protein A/G magnetic beads. CD25-interacting proteins were probed by Western blot analysis (**c**). Whole cell lysates (WCL) were used as positive controls for immunoblots. The isotype control for CD25 antibody (IgG) was used as negative control. EIF4E was used as specificity control. *Cd25*^fl/fl^ IL7-dependent pre-B cells, *BCR-ABL1-*transformed B-ALL, splenic mature-B cells and *Card11*^L232LI^-transformed mature B-cell lymphoma cells carrying 4-OHT-inducible Cre-ER^T2^ or ER^T2^ were treated with 4-OHT for 2 days (**d**). Cell lysates from these four cell types were studied by Western blot for protein levels of Ship1-pY^1020^, Ship1, Shp1-pY^564^, Shp1, Syk-pY^352^, Syk, Src-pY^416^, Lyn, Blnk-pY^96^, Blnk, Btk-pY^223^, Btk, PKCδ-pT^505^, PKCδ, Erk1/2-pT^202^/Y^204^ and Erk1/2. β-actin was used as loading control (**d**). Patient-derived B-ALL (**e**) cells (BCR-ABL1; PDX2) and TMD8 diffuse large B-cell lymphoma (DLBCL) cells (**f**) = were treated with Cas9 RNPs and guide RNAs for CRISPR-deletion of CD25 (gCD25) or non-targeting controls (gNT). PDX2 B-ALL cells were stimulated with the PKCδ-agonist ingenol-3-angelate (I3A) for an hour with different concentrations as indicated (**e**). IgM^+^ TMD8 DLBCL cells were stimulated with 1 µg/ml αIgM (F(ab’)_2_ fragments of anti-human μ chain) as indicated time points (**f**). Levels of Src-pY^416^, LYN, ERK1/2-pT^202^/Y^204^, ERK1/2, IκBα-pS^32/36^ and IκBα were assessed by Western blot, using β-actin as loading control. Murine *Cd25*^fl/fl^ *BCR-ABL1* B-ALL cells carrying an mClover Erk kinase translocation reporter (KTR) was transduced with 4-OHT-inducible Cre-ER^T2^ (Cre) or ER^T2^ (EV) and treated with 4-OHT for 2 days (**g**). In combination with Hoechst 33342 staining, Erk activity was assessed by the ratio of cytosolic to the nuclear intensity of mClover. Treatment with 1 µM PMA for 20 min was used as a positive control for nuclear Erk-translocation. Representative images are shown, scale bar, 5 µm. Statistical significance was determined by two-tailed Mann-Whitney *U*-test. *Cd25*^+/+^ *Mb1*-Cre-ER^T2^ and *Cd25*^fl/fl^ *Mb1*-Cre-ER^T2^ B-ALL cells (BCR-ABL1) were transduced with the GCaMP6s Ca^2+^ reporter to measure cytoplasmic Ca^2+^ mobilization (**h**). Representative time-course plots from single cells with a fold change of GCaMP6s intensity (y-axis) for a 3-s time interval (x-axis) in 2 min showed autonomous Ca^2+^ oscillations (middle). Fold changes of GCaMP6s intensity from 50 single cells are shown as heat-map (bottom). Representative images are shown (top). Scale bar, 100 µm. Ca^2+^ mobilization was assessed by GCaMP6s upon CRISPR-Cas9-mediated deletion of *CD25* with *CD25*-targeting guide RNAs (gCD25) or non-targeting guides (gNT) in patient-derived B-ALL (PDX2) cells (**i**). Representative time-course plots from single cell for 60 min with a 40-s time interval (x-axis) are shown (top). Fold changes of GCaMP6s intensity from 50 single cells are shown as heat-map (bottom). In **c-i**, Data are representative from at least three independent experiments. For gel source data, see **Supplementary Fig.1**.

## Phosphatase-activation depends on CD25-function in early and late stages of B-cell development

Since CD25-deletion had opposite effects on survival and proliferation in B-cell subsets and malignancies arising from early (pre-B cells, B-ALL) and late (mature B-cells, lymphoma) stages of B-cell development (**Figure 1d-e**), we examined the functional consequences of CD25-deletion on phosphatase-activation, PKCδ and BCR-downstream signaling in these four cellular contexts. Induced deletion of CD25 in murine pre-B, B-ALL, mature B- and B-cell lymphoma cells consistently resulted in loss of SHIP1- and SHP1-phosphorylation at sites reflecting phosphatase activation. In addition, the BCR-downstream kinases SYK and BTK and the linker BLNK as well as PKCδ were consistently hyperphosphorylated in all four cell types (**Figure 3d**). As in these mouse models, CRISPR-mediated deletion of CD25 in human B-ALL (early) and DLBCL (late) cells, consistently resulted in hyperactivation of BCR-downstream and NF-κB signaling pathways (**Figure 3e-f**).

## CD25 functions as negative feedback regulator of NF-κB signaling

Oncogenic activation of the NF-κB pathway by CARD11^L232LI^, IKK2^S177E^ and MYD88^L265P^ consistently induced transcriptional activation and surface expression of CD25 (**Extended Data Figure 1g, j-k**). Conversely, inducible deletion of CD25 resulted in increased activation of the NF-κB pathway, including PKCβ, RelA (p65) and IκBα (**Figure 3e-f; Extended Data Figure 4a**). These findings suggest a negative feedback loop in which NF-κB signaling induces transcriptional activation of CD25, which in turn assembles inhibitory phosphatases to attenuate kinase signaling, e.g., BTK and PKCβ (**Figure 3d, Extended Data Figure 4a**). CD25-deletion induced NF-κB activation in B-cells at both early (pre-B cells, B-ALL) and late (B-cells lymphoma) stages of development. Importantly, NF-κB activation has a tumor suppressive role in B-ALL and deletion of PKCβ and NF-κB1 accelerated development of B-ALL^48^. In contrast, oncogenic activation of NF-κB represents a central driver of mature B-cell lymphomas^42–44^. Taken together, these results suggest that stage-specific opposite effects of CD25-deletion reflect the known reverse outcomes of NF-κB activation in early^48^ and late stages^42–44^ of B-cell development. To assess whether increased NF-κB activation, at least in part, accounts for the divergent outcomes of CD25-deletion in B-ALL (early) and lymphoma (late) cells, we examined the MALT1 paracaspase small molecule inhibitor MI-2 and its ability to attenuate NF-κB signaling^49^. Interestingly, treatment with MI-2 was sufficient to mitigate toxicity of CD25-deletion in B-ALL cells. Conversely, CD25-deletion accelerated proliferation in mature B-cell lymphoma cells, which was reduced by MI-2 treatment. CD25-deletion amplified the proliferative response of mature B-cells to BCR-engagement, which was largely abrogated by MI-2 small molecule inhibition of NF-κB signaling (**Extended Data Figure 4b-d**).

## Divergent outcomes of CD25-deletion reflect stage-specific requirements of BCR-signaling strength

Biochemical studies in pre-B cells, B-ALL, mature B-cells, and lymphoma cells (**Figure 3d-e**) suggest that CD25 is uniformly required for the activation of inhibitory phosphatases and negative regulation of NF-κB, regardless of B-cell differentiation stage. Instead, the opposite outcomes of CD25-deletion during early and late stages of B-cell development are in line with distinct stage-specific requirements of regulation of BCR-signaling strength: Up to 75% of developing pre-B and immature B-cells are autoreactive^50^ and highly susceptible to negative selection by central tolerance mechanisms^51-53^. Owing to ubiquitous encounter of self-antigen, autoreactive B-cells exhibit hyperactive BCR-signaling, which induces anergy or clonal deletion^51-54^. Transitional (T1, T2) B-cells that arrive in the spleen still comprise about 40% autoreactive clones that remain highly susceptible to hyperactive BCR-signaling^52–54^. Interestingly, B-ALL^13,51,55–57^ and CLL^57,58^ cells that are derived from early and pre-GC stages of B-cell development share the same sensitivity to hyperactivation of BCR-signaling with their normal counterparts. In contrast, mature (follicular and marginal zone) and post-GC B-cells that have passed checkpoints for removal of autoreactive cells, are much less sensitive to hyperactive BCR-signaling^59–62^. Instead, vigorous BCR-signaling and NF-κB-activation promotes positive selection, survival, and proliferation at later stages of B-cell development. Likewise, chronic active^63^ and even autoreactive^64^ BCR-signaling function as a central oncogenic driver in B-cell lymphomas derived from post-GC B-cells. Hence, we propose that CD25 functions as a critical phosphatase-scaffold, which enables early B-cells, B-ALL and CLL cells to curb BCR-signaling and avoid negative selection during early B-cell development. At later stages of B-cell differentiation that are less sensitive to negative selection and central tolerance mechanisms, inhibitory CD25 signaling provides no competitive advantage and slows B-cell activation and proliferation of lymphoma cells.

## CD25 is required to prevent excessive activation of Erk, autonomous Ca^2+^ oscillations and B-cell anergy

Since hyperactive ERK-Ca^2+^-signaling in autoreactive transitional B-cells triggers negative selection and cell death^53–57^, we studied how genetic ablation of CD25 affected this pathway. Measuring activation of Erk by Western blot and an Erk-kinase translocation reporter (KTR), genetic deletion of CD25 caused hyperactivation of Erk (**Figure 3d-g**). Consistent with constitutive Erk and PKCδ-activation in CD25-deficient B-cells, Cre-mediated deletion of CD25 in murine B-ALL cells resulted in autonomous Ca^2+^-oscillations suggesting loss of negative regulation of this pathway in the absence of CD25 (**Figure 3h**). While patient-derived B-ALL cells showed occasional Ca^2+^-oscillations, their frequency was increased by ∼10-fold upon genetic deletion of CD25 (**Figure 3i**). Previous studies showed that hyperactivation of Erk and Ca^2+^-signaling in autoreactive B-cells results in anergy and negative selection during early B-cell development^53–55^. Consistent with these results, hyperactivation of the Erk-Ca^2+^ pathway upon loss of CD25 was associated with an NFAT- and anergy-related gene expression profile^65–67^ in pre-B cells and B-ALL, including upregulation of Nr4a1, Nr4a3, Egr2, Egr3, Ccl3, Batf, Maff and downregulation of Myc (**Extended Data Figure 5**). Consistent with previous work showing that mature B-cells are less sensitive to hyperactive BCR-signaling^59–62^, we found that activation of NFAT- and anergy-related gene expression programs upon CD25-deletion was reduced in mature B-cells and even reversed in *Card11*^L232LI^-driven B-cell lymphoma (**Extended Data Figure 5a-b**). A non-productive CD25 (Il2ra) transcript was among the most prominently upregulated mRNAs upon CD25-deletion in all four cell types. RNA-seq analysis revealed loss of exons 2-3, consistent with efficient Cre-mediated excision. Interestingly, however, loss of CD25 function, resulting in excessive BCR- and NF-κB activation, induced massive, albeit futile, transcriptional activation of CD25 as a negative feedback regulator, which is reflected by sharply increased RNA-seq counts of the non-functional CD25 transcript (**Extended Data Figure 5c**).

## Regulation of B-cell anergy by CD25

Induction of NFAT- and anergy-related gene expression programs represent central features of negative B-cell selection^65-67^. To directly study the role of CD25 in B-cell anergy, we studied B-cell-specific deletion of CD25 in double-transgenic mice expressing the neo-self-antigen hen egg lysozyme (HEL) and a high-affinity anti-HEL BCR (Ig^HEL^)^68^. In this model, Ig^HEL^ expressing B-cells are autoreactive and rapidly become anergic in the presence of soluble HEL (sHEL). Inducible deletion of CD25 in Ig^HEL^ B-cells exacerbated nuclear accumulation of NFAT and loss of BCR-responsiveness (**Extended Data Figure 6a-b**) as a functional hallmark of anergy^65–68^. As a direct readout of negative selection, inducible deletion of CD25 resulted in profound depletion of transitional Ig^HEL^ B-cells in the presence of sHEL (**Extended Data Figure 6c**). In addition, loss of CD25 resulted in upregulation of IgD at the expense of IgM and accumulation of transitional B-cells in the T3 compartment, which further corroborates a central role of CD25 in enabling immature and transitional B-cells to evade B-cell anergy and negative selection (**Extended Data Figure 6d**).

## CD25 is required to maintain B-cell anergy

Previous work demonstrated that continuous activity of the SHIP1 and SHP1 phosphatases is required to maintain unresponsiveness of anergic B-cells^69^. Since our biochemical studies demonstrated that CD25 is essential to activate inhibitory SHIP1 and SHP1 phosphatases (**Figure 3a-d**), we reasoned that deletion of CD25 and loss of SHIP1/SHP1-function might subvert B-cell anergy and allow anergic B-cells to regain responsiveness. To assess this idea, we cultured non-anergic Cd25^+/+^ Ig^HEL^, anergic Cd25^+/+^ Ig^HEL^ and anergic Cd25^-/-^ Ig^HEL^ B-cells. Non-anergic Cd25^+/+^ Ig^HEL^ B-cells (no sHEL) responded to stimulation with sHEL alone or in combination with IL4 or CD40L, entered cell cycle and proliferated with a low percentage of apoptotic cells. In contrast, anergic Cd25^+/+^ Ig^HEL^ B-cells (sHEL) remained unresponsive after restimulation and underwent apoptosis. Interestingly, anergic Cd25^-/-^ Ig^HEL^ B-cells (sHEL) not only regained BCR-responsiveness after restimulation and even exceeded responses of non-anergic B-cells with substantially increased proliferation and survival (**Extended Data Figure 6e-g**). Consistent with the critical role of inhibitory SHIP1 and SHP1 phosphatases in maintaining B-cell anergy^69^, these results highlight the functional importance of CD25 in the recruitment and activation of SHIP1 and SHP1 (**Figure 3a-d**).

## CD25 expressed on B-cells does not function as an IL2 receptor chain

Because CD25 was previously known as IL2-receptor chain on T- and NK-cells^6–9^, these results prompted us to investigate a previously unrecognized role of IL2-signaling in regulating inhibitory phosphatase activity. If the observed effects of CD25 on BCR-signaling and B-cell selection could be attributed to its canonical function in IL2 signaling in T- and NK-cells, one would expect that loss of IL2-signaling in *Il2*^-/-^ B-cells would phenocopy CD25-deficiency. However, the development and proportions of *Il2*^-/-^ B-cell subsets were normal and did not recapitulate the phenotypes of CD25-deletion (**Extended Data Figure 7a-b**). Rather than loss of IL2, CD25-deletion resembled phenotypes of genetic ablation of PKCδ^10–12^, SHIP1^13,14^ and SHP1^14-16^.

## CD25 interacts with the BCR but not IL2 receptor chains in B-cells

While in human peripheral blood T-cells and T-cell lymphoma cells, CD25 is coexpressed with the β- and γ-chains of the IL2 receptor^7–9^, we found that this is not the case in normal and transformed B-cells (**Extended Data Figure 7c-d**). Consistent with distinct functional roles, CD25 and the other chains of the IL2-receptor have different fates after endocytosis: CD25 has a half-life of >40 hours and is targeted to early endosomes for recycling to the cell membrane, whereas IL2Rβ and IL2Rγ chains with a one-hour half-life are targeted to late endosomes for degradation^34-35^. Proximity-ligation assays (PLA) in T-cell lymphoma cells showed ligand-dependent interactions of CD25 with IL2Rγ upon IL2-addition and with the TCR upon TCR-engagement. In B-cell lymphoma, however, CD25 only bound the BCR-signaling chain CD79B but not IL2Rγ (**Extended Data Figure 7c-e**). CD25:CD79B interactions in human B-cell lymphoma were markedly enhanced upon BCR-engagement, while IL2-treatment failed to induce CD25:IL2Rγ interactions (**Extended Data Figure 7c-e**). The PLA result for CD25:CD79B interactions in human B-cell lymphoma were corroborated by co-IP experiments in murine B-ALL cells (**Figure 3c**). These findings suggest a bivalent function of CD25 in IL2- and TCR-signaling in T-cells. However, in B-cells, CD25 is primarily associated with the BCR and does not contribute to IL2 signaling.

## IL2Rβ is required for CD25 recruitment to the IL2 receptor

Unlike CD25, IL2Rβ is essential for IL2-signal transduction in T-cells^7^. In contrast to IL2Rβ^+^ T-cell leukemia and lymphoma cells, normal and transformed B-cells lacked IL2Rβ expression and were not responsive to IL2 (**Extended Data Figure 8a-b**). Since expression of IL2Rβ was undetectable in B-cell malignancies, we induced ectopic expression of IL2Rβ in human B-cell lymphoma (JEKO1) cells (**Extended Data Figure 8c-e**). Ectopic expression of IL2Rβ in B-cell lymphoma cells was sufficient to enable IL2-signaling and CD25:IL2Rγ interactions in response to IL2. While BCR-engagement still resulted in CD25-recruitment to the CD79B signaling-chain of the BCR, IL2-mediated recruitment of CD25 to IL2Rγ was reduced when the BCR was concurrently engaged. Likewise, BCR-induced CD25:CD79B interactions were diminished when IL2 was concurrently added. These results suggest that BCR/TCR and the IL2-receptor can compete for recruitment of CD25 based on ligand-occupancy. While B-cells lack expression of a functional IL2 receptor, this model may be relevant to T-cells where CD25 could function as a bivalent coreceptor and interact with both TCR and IL2 receptor chains. In B-cell lymphoma cells that were engineered to express a functional IL2Rβ chain, IL2-mediated sequestration of CD25 from the BCR to IL2Rγ accelerated and increased Ca^2+^ signaling strength upon BCR-engagement (**Extended Data Figure 8e-f**). Consistent with a role of CD25 as negative regulator of BCR-signaling (**Figure 3**), IL2-induced sequestration of CD25 to IL2 receptor chains relieved inhibition of BCR-signaling.

## Recruitment of CD25 mirrors distinct types of oncogenic BCR-signaling in B-cell lymphomas

Some B-cell lymphoma subtypes exhibit tonic BCR-signaling, as opposed to chronic active BCR-signaling in most ABC DLBCL^63^, while other B-cell lymphomas lack or do not depend on oncogenic activation of the BCR-signaling pathway. To test whether distinct contexts of oncogenic BCR-signaling in specific B-cell lymphoma entities affects recruitment and function of CD25, we performed PLA experiments for CD25 and CD79B in lymphoma cell lines with either inducible (JEKO1) and chronic active (TMD8) BCR signaling or OCI-LY3 cells that were not dependent on proximal BCR-signaling as determined by BTK-kinase inhibitor resistance. In unstimulated JEKO1 mantle cell lymphoma cells, PLA experiments for CD25 and CD79B showed significant interactions, which were potentiated upon BCR-engagement. Unstimulated TMD8 ABC-DLBCL cells with chronic active BCR signaling showed a similar level of CD25:CD79B interactions as JEKO1 cells at baseline. Unlike JEKO1 cells, however, BCR-engagement did not further increase CD25:CD79B interactions. Consistent with lack of proximal BCR-signaling in OCI-LY3 cells, CD25:CD79B interactions were minimal at baseline and did not increase upon BCR-engagement (**Extended Data Figure 9**). Consistent with a central role of CD25 in feedback control of BCR-signaling, these results suggest that interactions between CD25 and the BCR are largely determined by the quality and strength of oncogenic activation of the BCR-signaling pathway.

## A PKCδ-motif in its cytoplasmic tail is critical for CD25-mediated feedback control of BCR-signaling

To elucidate the mechanism by which CD25 exerts feedback control of BCR-signaling in activated and transformed B-cells, we examined the role of its 12-aa cytoplasmic tail, including its central serine-phosphorylation site (S^268^). *In vitro* kinase assays of 62 serine/threonine kinases for a 12-aa peptide corresponding to the cytoplasmic tail of CD25 (WQRRQRK<u>S</u>^268^RRTI) and the S268A mutant peptide identified multiple PKC family members, including PKCδ, as upstream kinases based on selective phosphorylation of CD25 over the S268A mutant tail (**Figure 4a-b**). Given that PKCδ was the only inhibitory PKC family member^10–12^ in this group, we used a PKCδ-selective agonist, ingenol 3-angelate (I3A)^70,71^, to examine PKCδ-dependent phosphorylation and cell surface expression of CD25. Treatment of patient-derived B-ALL cells with I3A induced PKCδ-activation, CD25-phosphorylation on S^268^, and accumulation of CD25 at the cell surface (**Figure 4c-d**). In JEKO1 B-cell lymphoma cells, the PKCδ-agonist I3A amplified interactions between CD25 and the BCR-signaling chain CD79B (**Extended Data Figure 7e, 9a**). Engineering doxycycline-inducible reconstitution of wildtype and S268A-CD25 in CD25^-/-^ JEKO1 B-cell lymphoma cells revealed that S268A-mutant CD25 can be expressed at the cell surface. However, compared to wildtype CD25, surface expression of S268A-mutant CD25 was short-lived after one pulse of doxycycline (**Figure 4e-f**). Given that CD25 chains recycle between endosomal compartments and the cell surface^34-35^, these results suggest that PKCδ-mediated phosphorylation of S^268^ may delay CD25 endocytosis and internalization. Interestingly, CRISPR-mediated deletion of PKCδ (*PRKCD*) in JEKO1 B-cell lymphoma cells not only abolished phosphorylation and surface expression of CD25 but also reduced intracellular CD25 protein expression levels, as confirmed by Western blot (**Figure 4g-i**). Consistent with previous work demonstrating BCR-mediated activation and phosphorylation of PKCδ^71^, here we showed that BCR-dependent phosphorylation and stabilization of CD25 is mediated by PKCδ. BCR-engagement in *PRKCD*^-/-^ JEKO1 B-cell lymphoma cells failed to elicit CD25-phosphorylation on S^268^ (**Figure 4i**), demonstrating that CD25-tail phosphorylation depends on PKCδ rather than other PKC family members that were able to phosphorylate CD25-S^268^ *in vitro* (**Figure 4a**). In addition, CD25 protein levels in *PRKCD*^-/-^ JEKO1 B-cell lymphoma cells were reduced by 5- to 7-fold, suggesting that PKCδ-mediated phosphorylation may not only reduce internalization from the cell surface but also increase CD25 protein stability. We previously showed that CD25 function is essential for the survival and colony formation of B-ALL cells (**Figure 1d, h**). Consistent with a critical role of PKCδ in phosphorylation and stabilization of CD25, *Prkcd*^-/-^ pre-B cells were resistant to BCR-ABL1-mediated transformation and failed to develop B-ALL and to form colonies (**Figure 4j-k**). Previous work showed that individuals with inherited *PRKCD* germline point mutations suffer from B-cell defects and systemic autoimmunity^72^ as do patients with inherited CD25-mutations^26,27^. B-cells from patients with *PRKCD* point mutations (**Extended Data Figure 10**) lacked CD25-surface expression and increased expression of B-cell activation markers, including CD69 (**Figure 4l**). These results suggest that PKCδ exerts negative regulation of BCR-signaling and B-cell activation through phosphorylation and stabilization of CD25.

**Figure 4:**
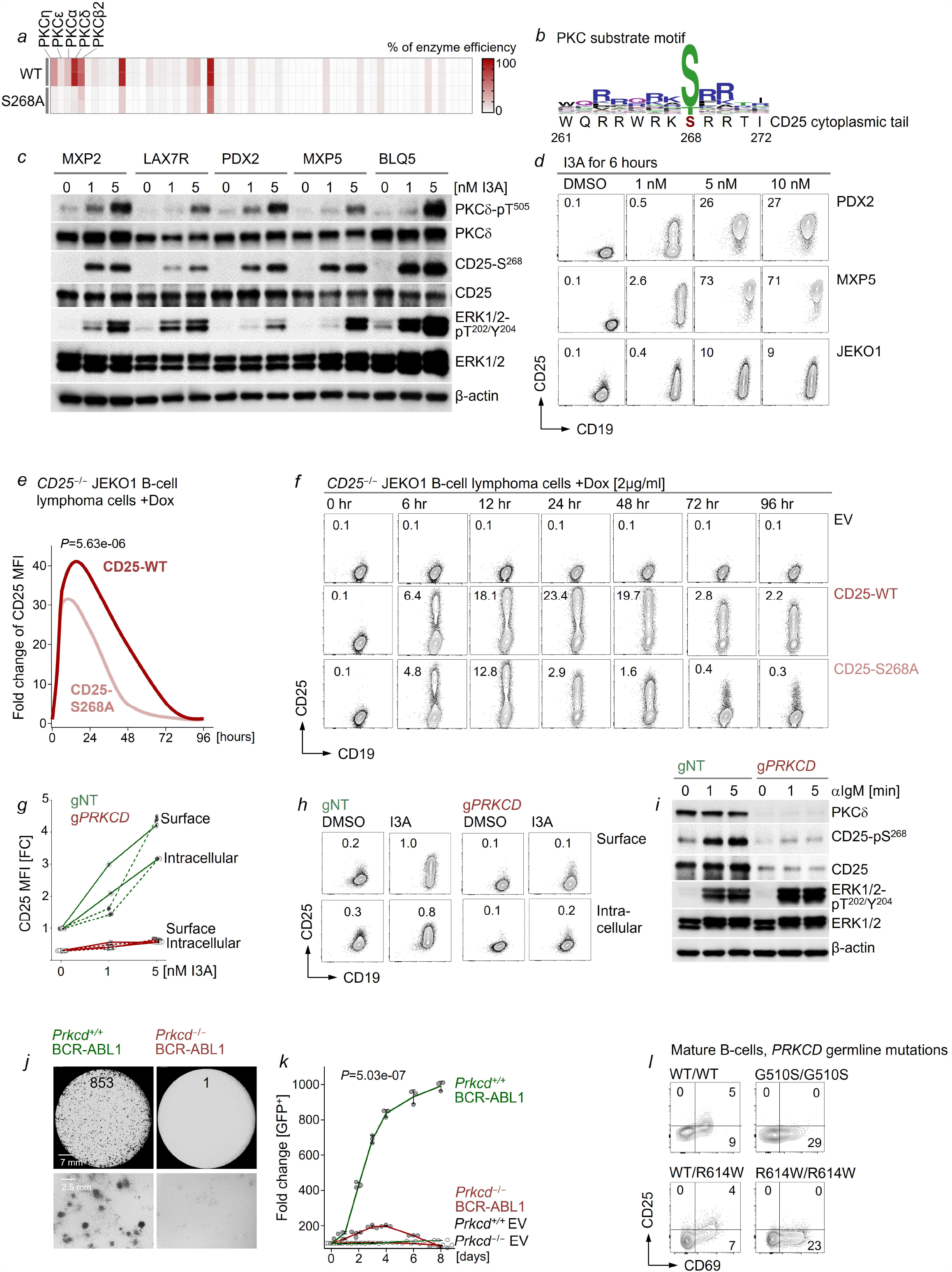
PKCδ-dependent phosphorylation of its cytoplasmic tail is required for CD25 cell surface expression. *In vitro* kinase assays of 62 known serine/threonine kinases against the recombinant cytoplasmic tail of CD25 (WQRRQRKS^268^RRTI) or S268A (WQRRQRKA^268^RRTI) were performed (**a**). The percentages of kinase efficiency quantitated as the percent of remaining ^33^P ATP (n=2) are represented as heatmap (see **Methods** and **Supplementary Table 1**). Schematic illustrating favored PKC substrate residues at each position on the cytoplasmic tail of CD25 (**b**). Western blot was performed to measure protein levels of PKCδ-pT^505^, PKCδ, CD25-pS^268^, CD25, ERK1/2-pT^202^/Y^204^, and ERK1/2 in five different patient-derived B-ALL cells (see **Supplementary Table 2**) upon treatment with the PKCδ-agonist I3A as indicated concentrations or vehicle (DMSO) for an hour. β-actin was used as loading control (**c**). Patient-derived B-ALL (PDX2; MXP5) and mantle cell lymphoma (JEKO1) cells treated with the indicated concentrations of I3A, or vehicle for 6 hours were analyzed by flow cytometry for surface expression of CD25 and CD19 (**d**). CD25 MFI (× 10^2^) values are indicated for individual measurements. *CD25*^-/-^ JEKO1 cells were reconstituted with doxycycline-inducible normal CD25 (CD25-WT) or S268A mutant (CD25-S268A; **e-f**). Levels of CD25 on surface were monitored by flow cytometry at indicated time points upon doxycycline induction and fold change of CD25 MFI was calculated (**e**). Representative plots of flow cytometric analysis are shown with CD25 MFI (× 10^2^) (**f**). PKCδ-deficient (*PRKCD*^-/-^) JEKO1 cells were generated by CRISPR-Cas9-mediated gene deletion (g*PRKCD*) and treated with I3A for six hours (dashed lines) or 12 hours (solid lines) at indicated concentrations. Combinations of surface and intracellular staining were conducted to examine CD25 expression and fold change of CD25 MFI was plotted (**g**). Representative plots of flow cytometric analysis are shown with CD25 MFI (× 10^2^). Non-targeting guide RNA (gNT) was used as control (**h**). PKCδ-deficient (gPRKCD) JEKO1 cells and non-targeting (gNT) controls were stimulated with anti-IgM antibodies for BCR-engagement. Cell lysates were probed for PKCδ to confirm deletion, CD25-pS^268^, CD25, ERK1/2-pT^202^/Y^204^, and ERK1/2 (**i**). β-actin was used as loading control. Murine *Prkcd*^+/+^ and *Prkcd*^-/-^ B-ALL cells transformed with BCR-ABL1 were plated in semi-solid methylcellulose for colony-forming assay (**j**). Representative images are shown with colony numbers. Scale bar, 7 mm (top row), 2.5 mm (bottom row). *Prkcd*^+/+^ and *Prkcd*^-/-^ pre-B cells were transduced with GFP-tagged BCR-ABL1 or empty vector (EV-GFP) in the presence of Imatinib (2 µmol/l). Enrichment or depletion of GFP^+^ cells upon acute activation of BCR-ABL1 by Imatinib washout on day 0 was monitored by flow cytometry (**k**) and fold changes of GFP^+^ populations were plotted (means ± s.d.). Expression of CD25 in primary B-cells isolated from peripheral blood in two patients with inherited *PRKCD* mutations and autoimmune disease and healthy siblings was assessed by flow cytometry after BCR-stimulation with 2 µg/ml anti-IgM for 2 days (**l**). Detailed information is included in **Extended Data Figure 10**. In **c-k**, Data are representative from at least three independent experiments. In **e, k**, Statistical significance was assessed by two-tailed *t*-test. For gel source data, see **Supplementary Fig.1**.

## Phosphorylation of the CD25 tail on S^268^ is required and sufficient for activation of inhibitory phosphatases

To study the mechanistic role of PKCδ-mediated phosphorylation of CD25-S^268^, we generated chimeric constructs containing the transmembrane domain (residues 239-261) and cytoplasmic tail (residues 262-272) of CD25 fused to the extracellular domain of CD19 (**Figure 5a**), a B-lymphoid transmembrane protein that is dispensable in B-ALL cells^22^. Consistent with dependency of B-ALL cells on CD25-function (**Figure 1d**), reconstitution of GFP-tagged CD25 expression conferred a substantial survival advantage in *CD25*^-/-^ B-ALL cells (**Figure 5b**). To elucidate structural requirements of CD25-mediated rescue, we compared constructs for full-length CD25, full-length CD19 and chimeric CD19:CD25 constructs. Expression of full-length CD19 had no significant effect on B-ALL cell survival and colony forming ability. Interestingly, the cytoplasmic tail of CD25 fused to CD19 was sufficient to rescue *CD25*^-/-^ B-ALL cells while the same construct carrying the S268A point mutation failed to confer a survival advantage and to restore colony formation capacity (**Figure 5c-d**). Likewise, the cytoplasmic tail of CD25, but not the S268A mutant, rescued activation of SHIP1 and SHP1 phosphatases and stabilized expression of the PKCδ scaffold RACK1 (**Figure 5e**). These results indicates that the PKCδ-substrate S^268^ in the tail of CD25 is required and sufficient to rescue survival, colony formation and activation of SHIP1 and SHP1 phosphatases in *CD25*^-/-^ B-ALL cells.

**Figure 5:**
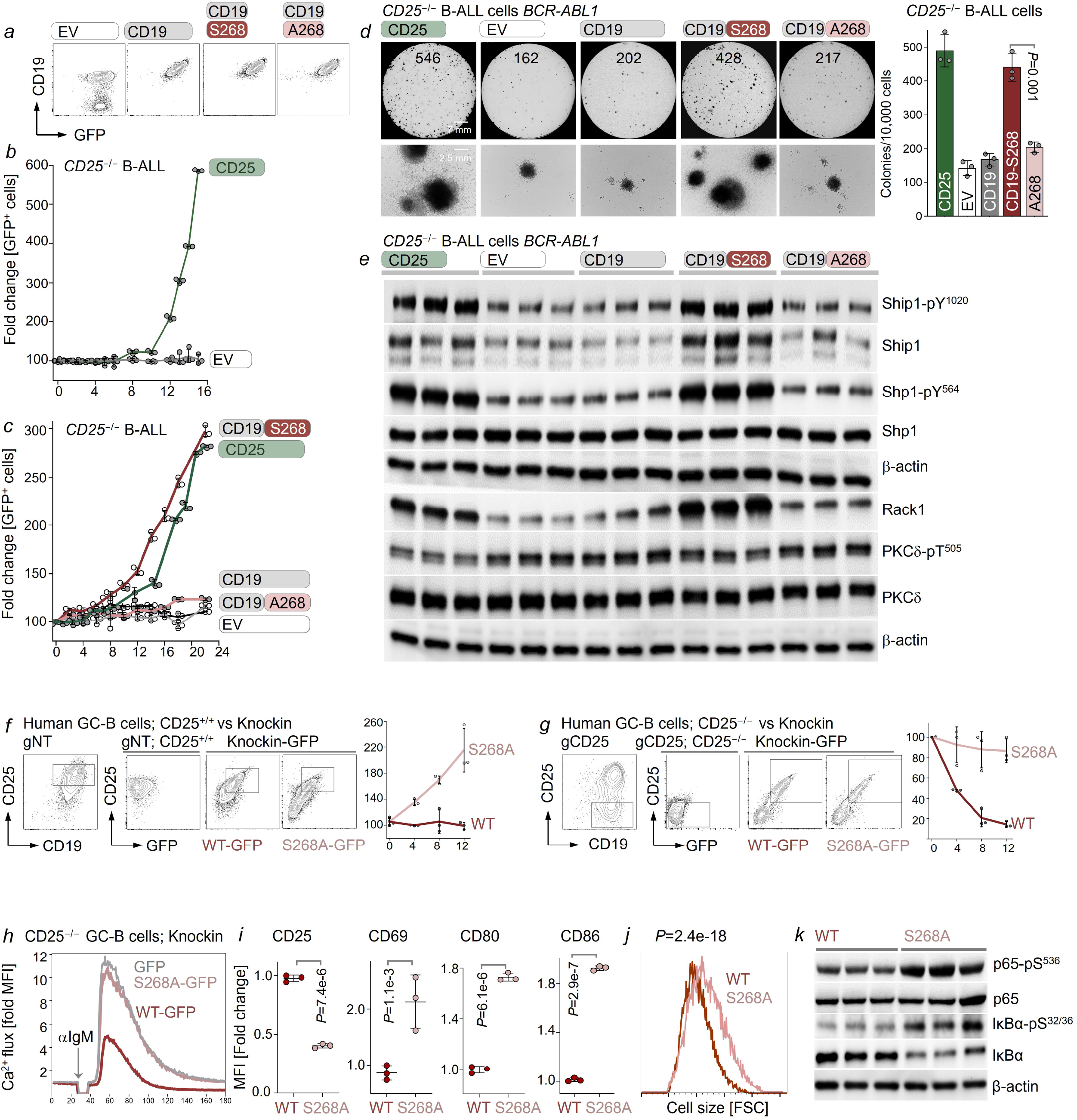
The PKCδ-phosphorylation motif of CD25 is required and sufficient for activation of inhibitory phosphatases in B-cells. Chimeric CD19:CD25 chains were constructed by fusing the extracellular domain of mouse CD19 (amino acids 1 to 287) to the transmembrane domain and cytoplasmic tail of the mouse CD25 (S^268^ or S268A) with GFP-tag. Empty vector (EV) or mouse CD19 (CD19) were used as control. Reconstitution of chimeric CD19:CD25 chains was validated by flow cytometry with detection of CD19 in combination with GFP expression in *Cd25*^-/-^ *BCR-ABL1* B-ALL cells (**a**). *Cd25*^-/-^ *BCR-ABL1* B-ALL cells were reconstituted with GFP-tagged wild-type CD25 or empty vector (EV). Enrichment of GFP^+^ cells was monitored by flow cytometry and fold change of GFP^+^ population (means ± s.d.) was plotted (**b**). *Cd25*^-/-^ *BCR-ABL1* B-ALL cells were reconstituted with GFP-tagged wild-type CD25, chimeric CD19:CD25 chains or empty vector (EV). Enrichment of GFP^+^ cells (means ± s.d.) was monitored by flow cytometry (**c**). *Cd25*^-/-^ *BCR-ABL1* B-ALL cells reconstituted with wild-type CD25, chimeric CD19:CD25 chains or empty vector (EV) were plated in semi-solid methylcellulose for colony-forming assays (**d**). Representative images are shown with colony numbers (means ± s.d.). Scale bar, 7 mm (top row), 2.5 mm (bottom row). Levels of Ship-pY^1020^, Ship, Shp1-pY^564^, Shp1, Rack1, PKCδ-pT^505^ and PKCδ were assessed by Western blot in *Cd25*^-/-^ *BCR-ABL1* B-ALL cells reconstituted with indicated constructs (**e**). β-actin was used as loading control. Human tonsillar germinal center (GC) B-cells enriched by negative selection were transduced with BCL2 and BCL6. Surface levels of CD25 were assessed following electroporation of non-targeting guide RNA (gNT) as control (**f**) or CD25-targeting guides (gCD25) for CRISPR-Cas9-mediated *CD25* deletion (**g**). CD25^+^ GC B-cells (gNT) and CD25^-^ GC B-cells (gCD25) were sorted by flow cytometry and subjected to growth competition assays (**f-g**). To this end, HDR templates encoding wild-type CD25 (S268)-GFP or S268A-GFP fusion protein were used for targeted knockin at the *CD25* locus in BCL2 and BCL6-transduced GC B-cells. GFP^+^ cells carrying the knockin alleles were sorted by flow cytometry and co-cultured with sorted CD25^+/+^ GFP^-^ cells (gNT; CD25^+/+^). Enrichment or depletion of GFP^+^ cells was monitored by flow cytometry and fold change of GFP^+^ population was plotted (means ± s.d.). Representative plots show levels of WT-GFP and S268A-GFP during growth competition assay (**f-g**). Ca^2+^ mobilization in response to BCR-engagement with F(ab’)_2_ fragments of anti-human μ chain was assessed for 180 sec in human CD25^-/-^ GC B-cells reconstituted with S268-GFP or S268A-GFP (**h**). Levels of CD25 and activation markers CD69, CD80 and CD86 were measured by flow cytometry (**i**) and relative MFIs were plotted (means ± s.d.). Levels of cell size measured by forward scatter were plotted (**j**). Human CD25^-/-^ GC B-cells were reconstituted with S268-GFP or S268A-GFP. GFP+ cells were sorted by flow cytometry and levels of NF-κB p65-pS^536^, NF-κB p65, IκBα-pS^32/36^, IκBα were accessed by Western blot, using β-actin as loading control (**k**). In **a-k**, Data are representative from at least three independent experiments. In **d, i, j**, two-tailed *t*-test. For gel source data, see **Supplementary Fig.1**.

## CD25-S^268^ enables negative feedback signaling in human germinal center B-cells

While deletion of CD25 had opposite outcomes in murine pre-B and mature B-cells (**Figure 1d-e**), we assessed if this was also the case for the S268A point mutation. To this end, we isolated primary human germinal center (GC) B-cells from tonsillectomy resectates, transduced GC B-cells with BCL2 and BCL6 to retain cell viability^73^ and induced CD25-deletion using non-targeting (NT) CRISPR-guides as control. In addition, we introduced genetic knockin alleles of CD25-GFP fusions using HDR templates^74^ encoding wildtype or S268A mutant CD25. To determine competitive fitness of GFP^+^ GC B-cells expressing CD25 wildtype or S268A knockin alleles, GFP^+^ cells were cultured in the presence of GFP^-^ CD25^+/+^ (NT-guides) or GFP^-^ CD25^-/-^ (CD25-targeting guides) GC B-cells. Fractions of GFP^+^ GC B-cells expressing the wildtype CD25-GFP knockin remained stable in competition with GFP^-^ CD25^+/+^ but were outcompeted by GFP^-^ CD25^-/-^ GC B-cells. Conversely, CD25-S268A knockin GC B-cells outcompeted their GFP^-^ CD25^+/+^ counterparts but remained on par with GFP^-^ CD25^-/-^ GC B-cells (**Figure 5f-g**). These results suggest that, unlike early stages of B-cell development, CD25 slows down proliferation of GC B-cells and that reduction of competitive fitness depends on the S^268^ site in its cytoplasmic tail. Genetic knockin of wildtype CD25-GFP in CD25^-/-^ GC B-cells reduced Ca^2+^ signaling upon BCR-engagement, whereas the CD25-S268A mutant, as well as the GFP control knock-in, failed to attenuate BCR-signaling (**Figure 5h**). Consistent with the requirement of PKCδ-mediated phosphorylation on S^268^ to stabilize CD25 (**Figure 4h-i**), protein levels of the CD25-S268A mutant were substantially lower level. In addition, CD25-S268A knockin GC B-cells expressed activation markers CD69, CD80 and CD86 at higher levels, had a larger cell size and exhibited constitutive activation of the NF-κB pathway (**Figure 5i-k**). While BCR-engagement and NF-κB-activation drive surface expression of CD25, these findings establish CD25-S^268^ and its interactions with PKCδ as a negative regulator of BCR-signaling and NF-κB activation in human GC B-cells.

## Structural basis of CD25-dependent recruitment and activation of inhibitory phosphatases

CD25-function and its ability to activate inhibitory SHIP1 and SHP1 phosphatases critically depend on the PKCδ-phosphorylation site S^268^. These findings are consistent with previous studies in mast cells and platelets suggesting that PKCδ plays an important role in activating inhibitory phosphatases^75,76^. Hence, we examined whether PKCδ-mediated phosphorylation of CD25-S^268^ provides the structural basis for recruitment of SHIP1 and SHIP phosphatases and other components of a CD25-dependent inhibitory complex. To investigate S^268^-dependent interactions of CD25, we performed Bio-ID proximity labeling experiments in human B-ALL (PDX2) and B-cell lymphoma (JEKO1) cells. CD25^-/-^ PDX2 and CD25^-/-^ JEKO1 cells were reconstituted with fusions between WT- or S268A-mutant CD25 and the biotin ligase BirA for labeling, streptavidin-pulldown, and mass spectrometry identification of interacting proteins. We used the CD25-S268A mutant as control for loss- and treatment with the PKCδ-agonist I3A^70^ and anti-IgM^71^ for gain- of PKCδ-mediated phosphorylation of CD25-S^268^ (**Figure 4c**). Comparing Bio-ID interactomes for CD25-WT and CD25-S268A identified SHIP1, SHP1 and the PKCδ-scaffold RACK1 among the CD25-interacting proteins that were most selective for CD25-WT over of the S268A mutant. Interestingly, treatment with the PKCδ-agonist I3A, and BCR-engagement in JEKO1 cells, intensified WT-selective interactions with the cytoplasmic tail of CD25 (**Figure 6a-b**). The dependency of interactions with SHIP1, SHP1 and the PKCδ-scaffold RACK1 on an intact CD25 cytoplasmic tail were validated by co-immunoprecipitation experiments. To this end, CD25-deficient PDX2 B-ALL and JEKO1 B-cell lymphoma cells were reconstituted with CD25-WT- or CD25-S268A. Western blot analyses using anti-CD25 antibodies confirmed that interactions between CD25 and SHIP1, SHP1 and RACK1 were largely dependent on S268 (**Figure 6c**). These results were further corroborated in proximity ligation assays (PLA) for SHP1 and RACK1 in JEKO1 B-cell lymphoma cells expressing CD25-WT- or CD25-S268A. Low baseline interactions between CD25-WT and SHP1 and RACK1 were dramatically increased by BCR-engagement. However, CD25-S268A lacked interactions with SHP1 and RACK1, regardless of BCR-engagement (**Figure 6d-e**).

**Figure 6:**
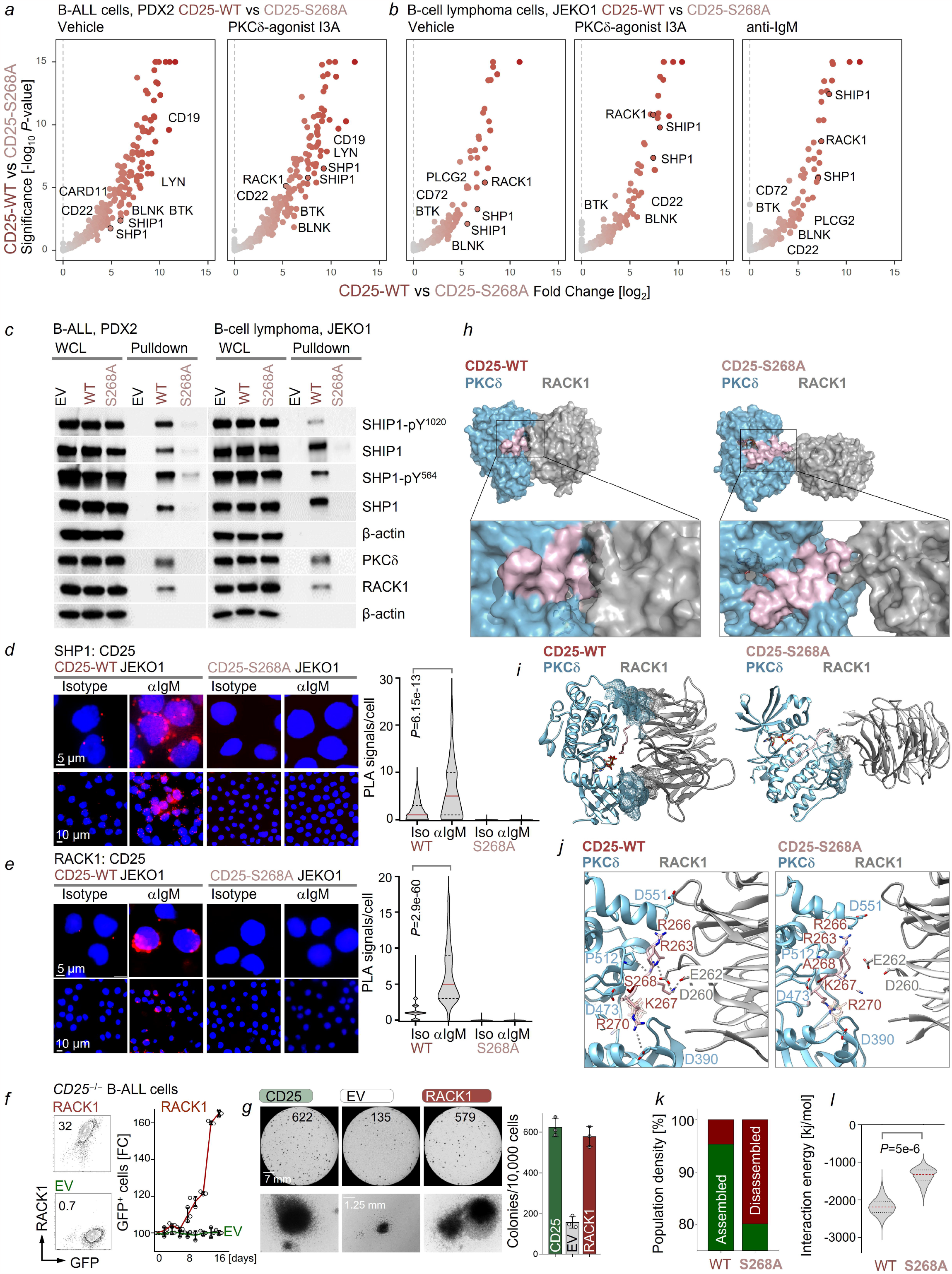
Structural basis of CD25-dependent recruitment and activation of inhibitory phosphatases. For Bio-ID interactome analyses, the BirA biotin ligase was fused to the C-terminus of wild-type (CD25-WT) and mutant (CD25-S268A) CD25. CD25^WT^-BirA, CD25^S268A^-BirA and BirA-EV control were reconstituted in *CD25*-deficient patient-derived B-ALL (PDX2) cells (**a**) or mature JEKO1 mantle cell lymphoma (MCL) cells (**b**). Cells were incubated with 50 μM exogenous biotin in culture media for 10 min upon treatment with the PKCδ-agonist I3A or BCR-engagement (JEKO1 cells only). To identify wild-type CD25-specific interactomes relative to CD25-S268A, log2-fold enrichment (x-axis) and significance (y-axis) of CD25^WT^-BirA vs CD25^S268A^-BirA in PDX2 and JEKO1 cells were plotted (n=4; moderated *t*-statistic). CD25^-/-^ PDX2 B-ALL or JEKO1 cells were reconstituted with wild-type (WT) or mutant (S268A) CD25, and co-immunoprecipitation using anti-CD25 antibody followed by Western blotting was performed to validate CD25-interacting proteins from Bio-ID interactome analyses. WCL was used as positive control for immunoblots. β-actin was used as specificity control (**c**). CD25^-/-^ JEKO1 cells were reconstituted with wild-type (WT) or mutant (S268A) CD25. Proximity ligation assays (PLA) were performed with reconstituted JEKO1 cells upon BCR-engagement. JEKO1 cells were stimulated with 10 μg/ml of F(ab’)_2_ fragments of anti-human μ chain or isotype control for 3 minutes, then fixed, permeabilized and assessed for proximity of SHP1 (**d**) or the PKCδ-scaffold RACK1 (**e**) to phosphorylated CD25-S^268^. Representative microscopic images with PLA signal (red dot) and nuclei stained with DAPI as blue are shown. IgG and CD25^-/-^ JEKO1 cells were used as negative controls. Scale bar, 5 µm (top row), 10 µm (bottom row). Quantitative analysis (right) shown as PLA signals per cell of each sample with the mean values shown as red line. *Cd25*^-/-^ *BCR-ABL1* B-ALL cells were reconstituted with constitutively active (myristoylated) RACK1-GFP or empty vector (EV; **f**). Expression levels of the PKCδ-scaffold RACK1 were validated by intracellular flow cytometry (left). Enrichment of GFP^+^ cells was monitored by flow cytometry (right; means ± s.d.). *Cd25*^-/-^ *BCR-ABL1* B-ALL cells reconstituted with wildtype CD25, EV or constitutively active (myristoylated) RACK1 were plated in semi-solid methylcellulose for colony-forming assays (**g**). Representative images are shown with colony numbers (means ± s.d.). Scale bar, 7 mm (top row), 2.5 mm (bottom row). Representative structures are shown as surface representation for the stable wild-type (CD25-WT; *left*) and less stable mutant CD25 (CD25-S268A; *right*) ternary complexes with PKCδ and RACK1. PKCδ is colored in blue, the cytoplasmic tail of CD25 is colored in pink and RACK1 is colored in gray (**h-j**). Same representative structures are shown in cartoon rendering for the ternary complex and ATP is shown in sticks. The blue and gray mesh regions denote contact regions in PKCδ and RACK1, respectively (**i**). Detailed PKCδ-RACK1 interaction interface for stable states (*left*) and initial step of weakening of the interface interactions transitioning to less stable states (*right*). Dashed lines label all the contacts which were persistent in over 60% of the simulation time. The residues were labeled in same color as the proteins they belong to (**j**). The ratios of population densities of stable (Assembled) versus less stable conformational states (Disassembled) in MD simulations of CD25-WT- or CD25-S268A-PKCδ-RACK1 ternary complexes were plotted (**k**). A cutoff of 6 Å in the root mean square deviations (RMSD) was applied to distinguish the two states (**k**). Combined interaction energy between PKCδ-CD25, CD25-RACK1 and PKCδ-RACK1 in CD25-WT- or CD25-S268A-PKCδ-RACK1 ternary complexes is shown as violin plot (**l**). In **c-g**, Data are representative from at least three independent experiments. In **a, b**, moderated *t*-test. In **d, e, i**, two-tailed *t*-test. For gel source data, see **Supplementary Fig.1**.

## Activation of the PKCδ-scaffold RACK1 can rescue loss of CD25-function

Previous work suggested that upon PKCδ-mediated phosphorylation, RACK1 can recruit inhibitory phosphatases and induce their activation^77–79^. As a result of these interactions, RACK1 functions as an inhibitory scaffold to negatively regulate Src and other tyrosine kinases^80–82^. For this reason, we determined whether constitutively active (myristoylated) RACK1 can functionally compensate for loss of CD25-function in B-ALL cells. Activated RACK1 restored competitive fitness and colony formation capability of CD25^-/-^ B-ALL cells to similar levels as reconstitution of CD25 (**Figure 6f-g**). These results suggests that CD25 orchestrates negative feedback regulation and recruitment of inhibitory phosphatases in B-cells by facilitating interactions between PKCδ and RACK1.

## Structural modeling of RACK1:CD25:PKCδ complexes

The initial three-dimensional structural model of the ternary RACK1:CD25:PKCδ complex model was constructed based on known features of the interaction between PKC isoforms and the blades 3 and 6 of RACK1^83–86^. In the absence of a crystal structure of full length PKCδ, we first modeled the catalytic domain of PKCδ with SWISS-model^87^, using PKCθ (PDB ID 5F9E) as template, and then modeled the substrate CD25-tail bound to PKCδ based on our previously validated structural model of substrate-bound PKCα^88^. The structural model of RACK1 was generated using homology modeling method with RACK1 structure as the template (PDB ID, 4OAW)^89^. The initial model of RACK1:CD25:PKCδ complex was energy minimized using Rosetta^90^ followed by 5 runs of all-atom molecular dynamics (MD) simulations^91^ (for details see **Methods section**). MD simulations of CD25^WT^ and CD25^S268A^-mutant with ATP and Mg^2+^ revealed two possible states of the RACK1:CD25:PKCδ complex. The RACK1:CD25^WT^:PKCδ complex favored a stable state. In contrast, RACK1:CD25^S268A^:PKCδ transitioned to a less stable complex with RACK1 departing from the PKCδ:CD25^S268A^ interface (**Figure 6h**). CD25^S268A^ retained some interactions with PKCδ, however contacts between RACK1 and CD25^S268A^ and PKCδ (van der Waals, hydrogen bond and salt-bridge) were almost entirely disrupted (**Figure 6i-j**; **Extended Data Figure 11a-b**). Calculation of conformational population densities of stable (Assembled) and unstable (Disassembled) states in MD simulation revealed that complexes with CD25^WT^ favored stable states whereas complexes with CD25^S268A^ favored less stable states (**Figure 6k**). The interaction energy for RACK1:CD25, RACK1:PKCδ and PKCδ:CD25 were calculated using GROMACS energy module^92^, and the sum of these three energies was used to evaluate the stability of the complex during MD simulation. The RACK1:CD25^WT^:PKCδ complex explored one major conformation in which the complex was stable with an interaction energy average at -2174 kJ/mol. Meanwhile, RACK1:CD25^S268A^:PKCδ formed a destabilized complex with an average interaction energy of - 1360 kJ/mol (**Figure 6l**), suggesting that the cytoplasmic tail of CD25 and PKCδ-mediated phosphorylation on S268 are required to form a stable complex.

## CD25 as a feedback regulator of oncogenic RTK signaling in acute myeloid leukemia

Previous work showed that in addition to activated B- and T-cells and lymphoid malignancies, CD25 can be expressed on a small subset of acute myeloid leukemia (AML). CD25 expression in these cases is associated with the oncogenic FLT3^ITD^ receptor-type tyrosine kinase (RTK) and particularly poor outcomes and aggressive disease (**Extended Data Figure 12a-b**). To study the role of CD25 in myeloid leukemia, we modeled AML based on the oncogenic fusion of MLL-AF9 transcription factors and the oncogenic FLT3^ITD^ RTK. Inducible expression of CD25 increased colony formation of FLT3^ITD^ AML cells (**Extended Data Figure 12c**). While Cre-mediated ablation of CD25 had no effect on MLL-AF9-driven AML, inducible ablation of CD25 in FLT3^ITD^ AML cells compromised colony formation and induced cell death. Likewise, loss of CD25 had no bearing on signal transduction in MLL-AF9-driven AML but introduced major signaling imbalances in FLT3^ITD^ AML cells, including increased AKT and ERK activity (**Extended Data Figure 12d-f**). These findings suggest that CD25 can function as feedback regulator in other cellular contexts: in analogy to recruitment of SHIP1 and SHP1 phosphatases for feedback control of BCR-signaling in B-cells, CD25 might have similar functions in the feedback regulations of RTKs, like FLT3^ITD^ in AML cells.

## CD25 surface expression represents a therapeutic target in B-cell malignancies

CD25-dependent feedback control of oncogenic signaling is essential for B-cell malignancies that arise from early (e.g., B-ALL) but not late (e.g., DLBCL) stages of B-cell development. We examined therapeutic targeting of CD25-surface expression in B-ALL and mature B-cell lymphomas. To this end, we studied a recently developed pyrrolobenzodiazepine (PBD) dimer-containing antibody-drug conjugate (ADCT-301)^93^ directed against human CD25. In patient-derived xenograft models for refractory B-ALL (BCR-ABL1), mantle cell lymphoma and DLBCL, ADCT-301 significantly extended overall survival of transplant recipients (**Extended Data Figure 13a-d**). Importantly, in refractory B-ALL PDX models, two injections of ADCT-301 achieved complete remission and long-term survival of all transplant recipients. While ADCT-301 also prolonged survival in mature B-cell malignancies (mantle cell lymphoma and DLBCL PDX), all transplant recipients ultimately died. Hence, targeting CD25 is likely of therapeutic benefit for both B-ALL and mature B-cell lymphomas. However, consistent with our mechanistic results that CD25 is essential in B-ALL but not in mature B-cell lymphoma, these results also suggest that targeting of B-ALL with ADCT-301 may be more durable and achieve more profound responses than mature B-cell lymphomas.

Here we identified CD25 as a negative feedback regulator of (pre-)BCR-signaling during normal B-cell development, as well as of oncogenes in B-cell malignancies that mimic BCR-downstream signaling pathways. While the β- and γ-chains of the IL2-receptor can be repurposed to become part of other cytokine receptors^37,38^, CD25 represents the first example of a cytokine receptor chain that can be recycled to recruit an inhibitory complex for feedback control of BCR-signaling or its oncogenic mimics. In conclusion, our findings highlight the previously unrecognized role of CD25 in assembling inhibitory phosphatases to control oncogenic signaling in B-cell malignancies and negative selection to prevent autoimmune disease.

## Acknowledgments

We would like to thank Lars Klemm and current and former members of the Müschen laboratory for their support and helpful discussions and Drs. Francesca Zammarchi and Patrick H. van Berkel (ADCT) for provision of CD25-ADC. Research in the Müschen laboratory is funded by the NIH through an NCI Outstanding Investigator Award R35CA197628, R01CA157644, R01CA213138, R01AI164692 and P01CA233412 (to M.M.), R01CA271497 (to J.C.), the Howard Hughes Medical Institute HHMI-55108547 (to M.M.), the Arthur H. and Isabel Bunker Chair in Hematology (to M.M.), a Blood Cancer Discovery Grant program through The Leukemia & Lymphoma Society, The Mark Foundation for Cancer Research, and The Paul G. Allen Frontiers Group and the V Foundation for Cancer Research T2018-003B (to M.M.).

M.M. is a Howard Hughes Medical Institute (HHMI) Faculty Scholar. J.L. was supported by Korea University Grant K2224161. V.S.V. was supported by the UCSF Grand Multiple Myeloma Translational Initiative (MMTI). Proteomics services were performed by the Northwestern Proteomics Core Facility, generously supported by NCI CCSG P30 CA060553 awarded to the Robert H Lurie Comprehensive Cancer Center, instrumentation award (S10OD025194) from NIH Office of Director, and the National Resource for Translational and Developmental Proteomics supported by P41 GM108569.

## References

1. Rolink, A., Grawunder, U., Winkler, T. H., Karasuyama, H. & Melchers, F. IL-2 receptor alpha chain (CD25, TAC) expression defines a crucial stage in pre-B cell development. Int Immunol 6, 1257–1264 (1994).

2. Muraguchi, A. et al. Interleukin 2 receptors on human B cells. Implications for the role of interleukin 2 in human B cell function. J Exp Med 161, 181 (1985).

3. Tumang, J. R. et al. c-Rel is essential for B lymphocyte survival and cell cycle progression. doi:10.1002/(SICI)1521-4141(199812)28:12.

4. Geng, H. et al. Integrative epigenomic analysis identifies biomarkers and therapeutic targets in adult B-acute lymphoblastic leukemia. Cancer Discov 2, 1006–1024 (2012).

5. Erber, W. N. & Mason, D. Y. Expression of the lnterleukin-2 Receptor (Tac Antigen/CD25) in Hematologic Neoplasms. Am J Clin Pathol 89, 645–648 (1988).

6. Leonard, W. J. et al. Molecular cloning and expression of cDNAs for the human interleukin-2 receptor. Nature 1984 311:5987 311, 626–631 (1984).

7. Nakamura, Y. et al. Heterodimerization of the IL-2 receptor β- and γ-chain cytoplasmic domains is required for signalling. Nature 1994 369:6478 369, 330–333 (1994).

8. Wang, X., Rickert, M. & Garcia, K. C. Structural biology: Structure of the quaternary complex of interleukin-2 with its α, β and γc receptors. Science (1979) 310, 1159–1163 (2005).

9. Stauber, D. J., Debler, E. W., Horton, P. A., Smith, K. A. & Wilson, I. A. Crystal structure of the IL-2 signaling complex: Paradigm for a heterotrimeric cytokine receptor. Proc Natl Acad Sci U S A 103, 2788–2793 (2006).

10. Mecklenbräuker, I., Saijo, K., Zheng, N. Y., Leitges, M. & Tarakhovsky, A. Protein kinase Cδ controls self-antigen-induced B-cell tolerance. Nature 2002 416:6883 416, 860–865 (2002).

11. Limnander, A. et al. Protein Kinase Cδ Promotes Transitional B Cell-Negative Selection and Limits Proximal B Cell Receptor Signaling to Enforce Tolerance. Mol Cell Biol 34, 1474–1485 (2014).

12. Miyamoto, A. et al. Increased proliferation of B cells and auto-immunity in mice lacking protein kinase Cδ. Nature 2002 416:6883 416, 865–869 (2002).

13. Ono, M., Bolland, S., Tempst, P. & Ravetch, J. v. Role of the inositol phosphatase SHIP in negative regulation of the immune system by the receptor FeγRIIB. Nature 1996 383:6597 383, 263–266 (1996).

14. Chen, Z. et al. Signalling thresholds and negative B-cell selection in acute lymphoblastic leukaemia. Nature 2015 521:7552 521, 357–361 (2015).

15. Cyster, J. G. & Goodnow, C. C. Protein tyrosine phosphatase 1C negatively regulates antigen receptor signaling in B lymphocytes and determines thresholds for negative selection. Immunity 2, 13–24 (1995).

16. Pao, L. I. et al. B Cell-Specific Deletion of Protein-Tyrosine Phosphatase Shp1 Promotes B-1a Cell Development and Causes Systemic Autoimmunity. Immunity 27, 35–48 (2007).

17. Reth, M. & Alarcon B Wileman, C. H. Antigen receptor tail clue. Nature 1989 338:6214 338, 383–384 (1989).

18. Vivier, E. & Daëron, M. Immunoreceptor tyrosine-based inhibition motifs. Immunol Today 18, 286–291 (1997).

19. Clemens, L. et al. Determination of the molecular reach of the protein tyrosine phosphatase SHP-1. Biophys J 120, 2054–2066 (2021).

20. Haslam, R. J., Koide, H. B. & Hemmings, B. A. Pleckstrin domain homology. Nature 1993 363:6427 363, 309–310 (1993).

21. Mayer, B. J., Ren, R., Clark, K. L. & Baltimore, D. A putative modular domain present in diverse signaling proteins. Cell 73, 629–630 (1993).

22. Lee, J. et al. IFITM3 functions as a PIP3 scaffold to amplify PI3K signalling in B cells. Nature 2020 588:7838 588, 491–497 (2020).

23. Teshigawara, K., Wang, H. M., Kato, K. & Smith, K. A. Interleukin 2 high-affinity receptor expression requires two distinct binding proteins. Journal of Experimental Medicine 165, 223–238 (1987).

24. Leonard, W. J. et al. Cytoplasmic domains of the interleukin-2 receptor β and γ chains mediate the signal for T-cell proliferation. Nature 1994 369:6478 369, 333–336 (1994).

25. Goudy, K. et al. Human IL2RA null mutation mediates immunodeficiency with lymphoproliferation and autoimmunity. Clinical Immunology 146, 248–261 (2013).

26. Sharfe, N., Dadi, H. K., Shahar, M. & Roifman, C. M. Human immune disorder arising from mutation of the α chain of the interleukin-2 receptor. Proc Natl Acad Sci U S A 94, 3168–3171 (1997).

27. Willerford, D. M. et al. Interleukin-2 receptor α chain regulates the size and content of the peripheral lymphoid compartment. Immunity 3, 521–530 (1995).

28. Lowe, C. E. et al. Large-scale genetic fine mapping and genotype-phenotype associations implicate polymorphism in the IL2RA region in type 1 diabetes. Nature Genetics 2007 39:9 39, 1074–1082 (2007).

29. Sawcer, S. et al. Genetic risk and a primary role for cell-mediated immune mechanisms in multiple sclerosis. Nature 2011 476:7359 476, 214–219 (2011).

30. Hafler, D. A. et al. Risk Alleles for Multiple Sclerosis Identified by a Genomewide Study. New England Journal of Medicine 357, 851–862 (2007).

31. Hinks, A. et al. Association of the IL2RA/CD25 gene with juvenile idiopathic arthritis. Arthritis Rheum 60, 251–257 (2009).

32. Carr, E. J. et al. Contrasting genetic association of IL2RA with SLE and ANCA - Associated vasculitis. BMC Med Genet 10, 1–7 (2009).

33. Chang, J. S. et al. Genetic polymorphisms in adaptive immunity genes and childhood acute lymphoblastic leukemia. Cancer Epidemiology Biomarkers and Prevention 19, 2152–2163 (2010).

34. Hémar, A. et al. Endocytosis of interleukin 2 receptors in human T lymphocytes: distinct intracellular localization and fate of the receptor alpha, beta, and gamma chains. Journal of Cell Biology 129, 55–64 (1995).

35. Hémar, A. & DautryLVarsat, A. Cyclosporin A inhibits the interleukin 2 receptor α chain gene transcription but not its cell surface expression: the α chain stability can explain this discrepancy. Eur J Immunol 20, 2629–2635 (1990).

36. Mutation Analysis of IL2RG in Human X-Linked Severe Combined Immunodeficiency. Blood 89, 1968–1977 (1997).

37. Zurawski, 21 S M et al. Interleukin-2 Receptor γ Chain: A Functional Component of the Interleukin-7 Receptor. Science (1979) 262, 1877–1880 (1993).

38. Grabstein, K. H. et al. Cloning of a T Cell Growth Factor that Interacts with the β Chain of the Interleukin-2 Receptor. Science (1979) 264, 965–968 (1994).

39. Geng, H. et al. Self-Enforcing Feedback Activation between BCL6 and Pre-B Cell Receptor Signaling Defines a Distinct Subtype of Acute Lymphoblastic Leukemia. Cancer Cell 27, 409–425 (2015).

40. Trageser, D. et al. Pre–B cell receptor–mediated cell cycle arrest in Philadelphia chromosome– positive acute lymphoblastic leukemia requires IKAROS function. J Exp Med 206, 1739 (2009).

41. Feldhahn, N. et al. Mimicry of a constitutively active pre–B cell receptor in acute lymphoblastic leukemia cells. Journal of Experimental Medicine 201, 1837–1852 (2005).

42. Lenz, G. et al. Oncogenic CARD11 mutations in human diffuse large B cell lymphoma. Science (1979) 319, 1676–1679 (2008).

43. Ngo, V. N. et al. Oncogenically active MYD88 mutations in human lymphoma. Nature 2010 470:7332 470, 115–119 (2010).

44. Eric Davis, R., Brown, K. D., Siebenlist, U. & Staudt, L. M. Constitutive Nuclear Factor κB Activity Is Required for Survival of Activated B Cell–like Diffuse Large B Cell Lymphoma Cells. Journal of Experimental Medicine 194, 1861–1874 (2001).

45. Chen, Y. et al. SHIP-1 Deficiency in AID+ B Cells Leads to the Impaired Function of B10 Cells with Spontaneous Autoimmunity. The Journal of Immunology 199, 3063–3073 (2017).

46. Sander, S. et al. PI3 Kinase and FOXO1 Transcription Factor Activity Differentially Control B Cells in the Germinal Center Light and Dark Zones. Immunity 43, 1075–1086 (2015).

47. Rameh, L. E., Tolias, K. F., Duckworth, B. C. & Cantley, L. C. A new pathway for synthesis of phosphatidylinositol-4,5-bisphosphate. Nature 1997 390:6656 390, 192–196 (1997).

48. Katerndahl, C. D. S. et al. Antagonism of B cell enhancer networks by STAT5 drives leukemia and poor patient survival. Nature Immunology 2017 18:6 18, 694–704 (2017).

49. Fontan, L. et al. MALT1 Small Molecule Inhibitors Specifically Suppress ABC-DLBCL In Vitro and In Vivo. Cancer Cell 22, 812–824 (2012).

50. Wardemann, H. et al. Predominant autoantibody production by early human B cell precursors. Science (1979) 301, 1374–1377 (2003).

51. Shojaee, S. et al. PTEN opposes negative selection and enables oncogenic transformation of pre-B cells. Nature Medicine 2016 22:4 22, 379–387 (2016).

52. Yurasov, S. et al. Defective B cell tolerance checkpoints in systemic lupus erythematosus. J Exp Med 201, 703–711 (2005).

53. Limnander, A. et al. STIM1, PKC-δ and RasGRP set a threshold for proapoptotic Erk signaling during B cell development. Nat Immunol 12, 425–433 (2011).

54. Zikherman, J., Parameswaran, R. & Weiss, A. Endogenous antigen tunes the responsiveness of naive B cells but not T cells. Nature 2012 489:7414 489, 160–164 (2012).

55. Shojaee, S. et al. Erk Negative Feedback Control Enables Pre-B Cell Transformation and Represents a Therapeutic Target in Acute Lymphoblastic Leukemia. Cancer Cell 28, 114–128 (2015).

56. Müschen, M. Autoimmunity checkpoints as therapeutic targets in B cell malignancies. Nature Reviews Cancer 2018 18:2 18, 103–116 (2018).

57. Sadras, T. et al. Developmental partitioning of SYK and ZAP70 prevents autoimmunity and cancer. Mol Cell 81, 2094-2111.e9 (2021).

58. Ecker, V. et al. Targeted PI3K/AKT-hyperactivation induces cell death in chronic lymphocytic leukemia. Nature Communications 2021 12:1 12, 1–17 (2021).

59. Su, T. T., Guo, B., Wei, B., Braun, J. & Rawlings, D. J. Signaling in transitional type 2 B cells is critical for peripheral B-cell development. Immunol Rev 197, 161–178 (2004).

60. Su, T. T. & Rawlings, D. J. Transitional B lymphocyte subsets operate as distinct checkpoints in murine splenic B cell development. J Immunol 168, 2101–2110 (2002).

61. Pillai, S. & Cariappa, A. The follicular versus marginal zone B lymphocyte cell fate decision. Nature Reviews Immunology 2009 9:11 9, 767–777 (2009).

62. Pillai, S., Cariappa, A. & Moran, S. T. Positive selection and lineage commitment during peripheral B-lymphocyte development. Immunol Rev 197, 206–218 (2004).

63. Davis RE, Ngo VN, Lenz G et al. Chronic active B-cell-receptor signalling in diffuse large B-cell lymphoma. Nature 463, 88–92 (2010).

64. Young RM, Wu T, Schmitz R et al. Survival of human lymphoma cells requires B-cell receptor engagement by self-antigens. Proc Natl Acad Sci U S A. 112, 13447–54 (2015).

65. Barrington, R. A., Borde, M., Rao, A. & Carroll, M. C. Involvement of NFAT1 in B Cell Self-Tolerance. The Journal of Immunology 177, 1510–1515 (2006).

66. Märklin, M. et al. NFAT2 is a critical regulator of the anergic phenotype in chronic lymphocytic leukaemia. Nature Communications 2017 8:1 8, 1–14 (2017).

67. Tan, C. et al. NR4A nuclear receptors restrain B cell responses to antigen when second signals are absent or limiting. Nature Immunology 2020 21:10 21, 1267–1279 (2020).

68. Goodnow, C. C. et al. Altered immunoglobulin expression and functional silencing of self-reactive B lymphocytes in transgenic mice. Nature 334, 676–682 (1988).

69. Getahun, A., Beavers, N. A., Larson, S. R., Shlomchik, M. J. & Cambier, J. C. Continuous inhibitory signaling by both SHP-1 and SHIP-1 pathways is required to maintain unresponsiveness of anergic B cells. J Exp Med 213, 751–769 (2016).

70. Kedei, N. et al. Characterization of the Interaction of Ingenol 3-Angelate with Protein Kinase C. Cancer Res 64, 3243–3255 (2004).

71. Pracht, C., Minguet, S., Leitges, M., Reth, M. & Huber, M. Association of protein kinase C-delta with the B cell antigen receptor complex. Cell Signal 19, 715–722 (2007).

72. Salzer, E. et al. B-cell deficiency and severe autoimmunity caused by deficiency of protein kinase C δ. Blood 121, 3112–3116 (2013).

73. Caeser, R. et al. Genetic modification of primary human B cells to model high-grade lymphoma. Nature Communications 2019 10:1 10, 1–16 (2019).

74. Shy, B. R. et al. High-yield genome engineering in primary cells using a hybrid ssDNA repair template and small-molecule cocktails. Nature Biotechnology 2022 1–11 (2022) doi:10.1038/s41587-022-01418-8.

75. Leitges, M. et al. Protein kinase C-delta is a negative regulator of antigen-induced mast cell degranulation. Mol Cell Biol 22, 3970–3980 (2002).

76. Chari, R. et al. Lyn, PKC-delta, SHIP-1 interactions regulate GPVI-mediated platelet-dense granule secretion. Blood 114, 3056–3063 (2009).

77. Rosdahl, J. A., Mourton, T. L. & Brady-Kalnay, S. M. Protein Kinase C δ (PKCδ) Is Required for Protein Tyrosine Phosphatase μ (PTPμ)-Dependent Neurite Outgrowth. Molecular and Cellular Neuroscience 19, 292–306 (2002).

78. Kiely, P. A., O’Gorman, D., Luong, K., Ron, D. & O’Connor, R. Insulin-like growth factor I controls a mutually exclusive association of RACK1 with protein phosphatase 2A and beta1 integrin to promote cell migration. Mol Cell Biol 26, 4041–4051 (2006).

79. Mourton, T. et al. The PTPmu protein-tyrosine phosphatase binds and recruits the scaffolding protein RACK1 to cell-cell contacts. J Biol Chem 276, 14896–14901 (2001).

80. Chang, B. Y., Conroy, K. B., Machleder, E. M. & Cartwright, C. A. RACK1, a receptor for activated C kinase and a homolog of the beta subunit of G proteins, inhibits activity of src tyrosine kinases and growth of NIH 3T3 cells. Mol Cell Biol 18, 3245–3256 (1998).

81. Mamidipudi, V. et al. RACK1 inhibits colonic cell growth by regulating Src activity at cell cycle checkpoints. Oncogene 2007 26:20 26, 2914–2924 (2006).

82. Miller, L. D., Lee, K. C., Mochly-Rosen, D. & Cartwright, C. A. RACK1 regulates Src-mediated Sam68 and p190RhoGAP signaling. Oncogene 2004 23:33 23, 5682–5686 (2004).

83. Ron, D. et al. Cloning of an intracellular receptor for protein kinase C: a homolog of the beta subunit of G proteins. Proc Natl Acad Sci U S A 91, 839–843 (1994).

84. Ron, D. & Mochly-Rosen, D. Agonists and antagonists of protein kinase C function, derived from its binding proteins. Journal of Biological Chemistry 269, 21395–21398 (1994).

85. Ron, D. & Mochly-Rosen, D. An autoregulatory region in protein kinase C: the pseudoanchoring site. Proc Natl Acad Sci U S A 92, 492 (1995).

86. Jones, A. C., Taylor, S. S., Newton, A. C. & Kornev, A. P. Hypothesis: Unifying model of domain architecture for conventional and novel protein kinase C isozymes. IUBMB Life 72, 2584–2590 (2020).

87. Waterhouse, A. et al. SWISS-MODEL: homology modelling of protein structures and complexes. Nucleic Acids Res 46, W296–W303 (2018).

88. Lee, S. et al. Distinct structural mechanisms determine substrate affinity and kinase activity of protein kinase Cα. J Biol Chem 292, 16300–16309 (2017).

89. Ruiz Carrillo, D. et al. Structure of human Rack1 protein at a resolution of 2.45 Å. Acta Crystallogr 68, 867–872 (2012).

90. Alford, R. F. et al. The Rosetta All-Atom Energy Function for Macromolecular Modeling and Design. J Chem Theory Comput 13, 3031–3048 (2017).

91. Huang, J. et al. CHARMM36m: an improved force field for folded and intrinsically disordered proteins. Nature Methods 2016 14:1 14, 71–73 (2016).

92. Berendsen, H. J. C., van der Spoel, D. & van Drunen, R. GROMACS: A message-passing parallel molecular dynamics implementation. Comput Phys Commun 91, 43–56 (1995).

93. Zammarchi, F. et al. CD25-targeted antibody-drug conjugate depletes regulatory T cells and eliminates established syngeneic tumors via antitumor immunity. J Immunother Cancer 8, (2020).

