## Extended Data Figures for "Dynamic phosphatase-recruitment controls B-cell selection and oncogenic signaling"

**Extended Data Figure 1:** *(pre-)BCR-signaling and its oncogenic mimics drive CD25 expression in B-cells*

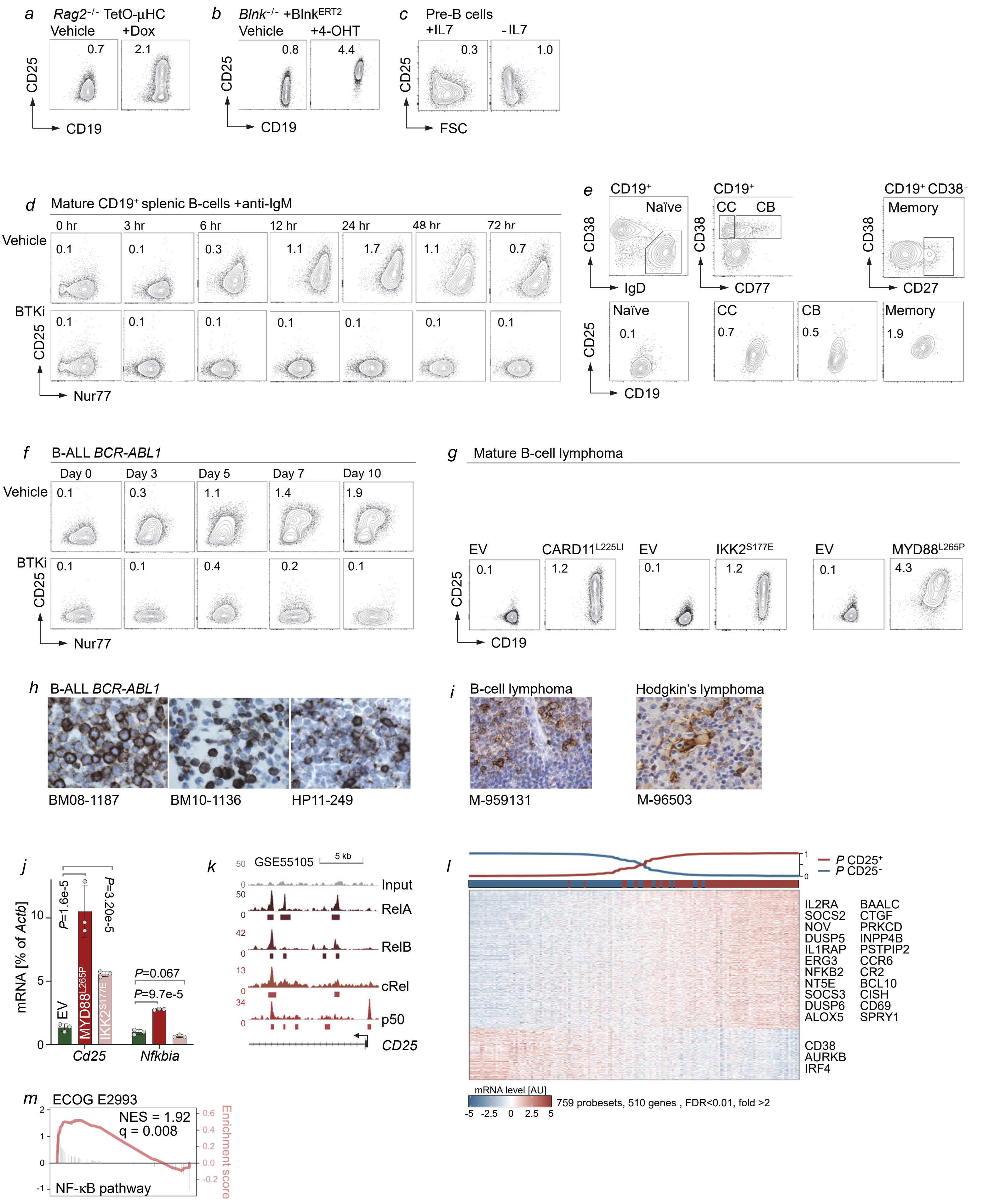

**Extended Data Figure 1:** *(pre-)BCR-signaling and its oncogenic mimics drive CD25 expression in B-cells*

**a-c,** Murine *Rag2*$^{-/-}$ Il-7-dependent pre-B cells were reconstituted with doxycycline (Dox)-inducible TetO-μHC and treated with vehicle or doxycycline for 24 hours. Expression of CD25 was measured by flow cytometry upon reconstitution of μHC (**a**).

Murine *Blnk*$^{-/-}$ Il-7-dependent pre-B cells were reconstituted with 4-OHT-inducible Blnk-ER$^{T2}$ encoding fusion protein of the ER-ligand binding domain with the N terminus of Blnk. Upregulation of CD25 upon release of Blnk-ER$^{T2}$ from cytoplasmic heatshock chaperone by treatment of 1 μM 4-OHT for 24 hours was measured by flow cytometry (**b**).

Levels of CD25 and forward scatter in normal Il-7-dependent pre-B cells were assessed after withdrawal of Il-7 for 3 days (**c**). MFIs (× $10^2$) for CD25 are indicated. **d,** Splenic B-cells purified by negative selection from Nur77-GFP mice were stimulated with 10 μg/ml anti-IgM for various lengths of time in the presence of vehicle control or Ibrutinib (10 μM). Representative dot plots of CD25 and Nur77-GFP expression gated on CD19$^+$ are shown. MFIs (× $10^2$) for CD25 are depicted.

**e,** Human tonsillar B-cells were purified by negative selection and CD25 expression was assessed in naïve B-cells (CD19$^+$, CD38$^{low}$, IgD$^+$, CD27$^-$), centrocytes (CC, CD19$^+$, CD38$^{high}$, CD77$^-$), centroblasts (CB, CD19$^+$, CD38$^{high}$, CD77$^+$) and memory B cells (CD19$^+$, CD38$^{low}$, CD27$^+$). MFIs (× $10^3$) for CD25 are indicated. **f,** Murine IL-7-dependent pre-B cells from Nur77-GFP mice were transduced with BCR-ABL1 on day 0 and expression of CD25 and Nur77-GFP was monitored by flow cytometry at different time points in the presence of vehicle control or Ibrutinib (10 μM). MFIs (× $10^2$) for CD25 are depicted.

**g,** Human tonsillar B-cells purified by negative selection were transduced with empty vector (EV), CARD11$^{L225LI}$, IKK2$^{S177E}$, MYD88$^{L265P}$. Expression of CD25 was measured and MFIs (× $10^3$) for CD25 are indicated. **h-i,** Immunohistochemistry staining for CD25 (brown) of paraffin-embedded bone marrow samples (n=3) from *BCR-ABL1* (**h**) or lymph node (n=2) from Non-Hodgkin's B cell lymphoma (**i**, left) and Hodgkin's lymphoma **i**, right).

**j,** Murine IL-7-dependent pre-B cells were transduced with MYD88$^{L265P}$, IKK2$^{S177E}$ or empty vector (EV) control and analyzed for *Cd25* and *Nfkbia* mRNA levels by quantitative RT-PCR relative to *Actb* (n=3).

**k,** ChIP-seq analysis for binding of NF-κB subunits RelA, RelB, cRel and p50 at the *CD25* locus in human lymphoblastoid B-cells, GM12878 (GSE55105), bars indicate significant peaks. **l,** Gene expression microarray data of B-ALL patient samples from ECOG E2993 (GEO: GSE34861) identified 510 genes (759 probesets) were significantly up- or down-regulated ($\log_2$ fold change >2, FDR <0.01) in CD25$^+$ (defined as >20% positive cells by flow cytometry) versus CD25$^-$ samples (defined as <1% positive cells). Adjusted *P*-values were calculated by Student *t*-test with Benjamini–Hochberg correction for multiple testing. The probability of a Bayesian predictor to reclassify CD25$^+$ (*P* CD25$^+$) or CD25$^-$ (*P* CD25$^-$) ALL cases are shown on top of the heatmap. The number of correctly classified or misclassified cases revealed a classification accuracy of 94%. **m,** Gene set enrichment analysis showed a significant enrichment of the NF-κB pathway in CD25$^+$ versus CD25$^-$ patient enriched genes from ECOG E2993. Red lines indicate running enrichment score (right-axis), grey bars indicate log2 fold change (left-axis).

Statistical significance was determined by two-tailed Kolmogorov-Smirnov test. In **a-g, j,** Data are representative from at least three independent experiments. In **j,** two-tailed *t*-test (means ± s.d.).

**Extended Data Figure 2:** *CD25 functions as a feedback regulator and promotes drug-resistance in patient-derived B-ALL*

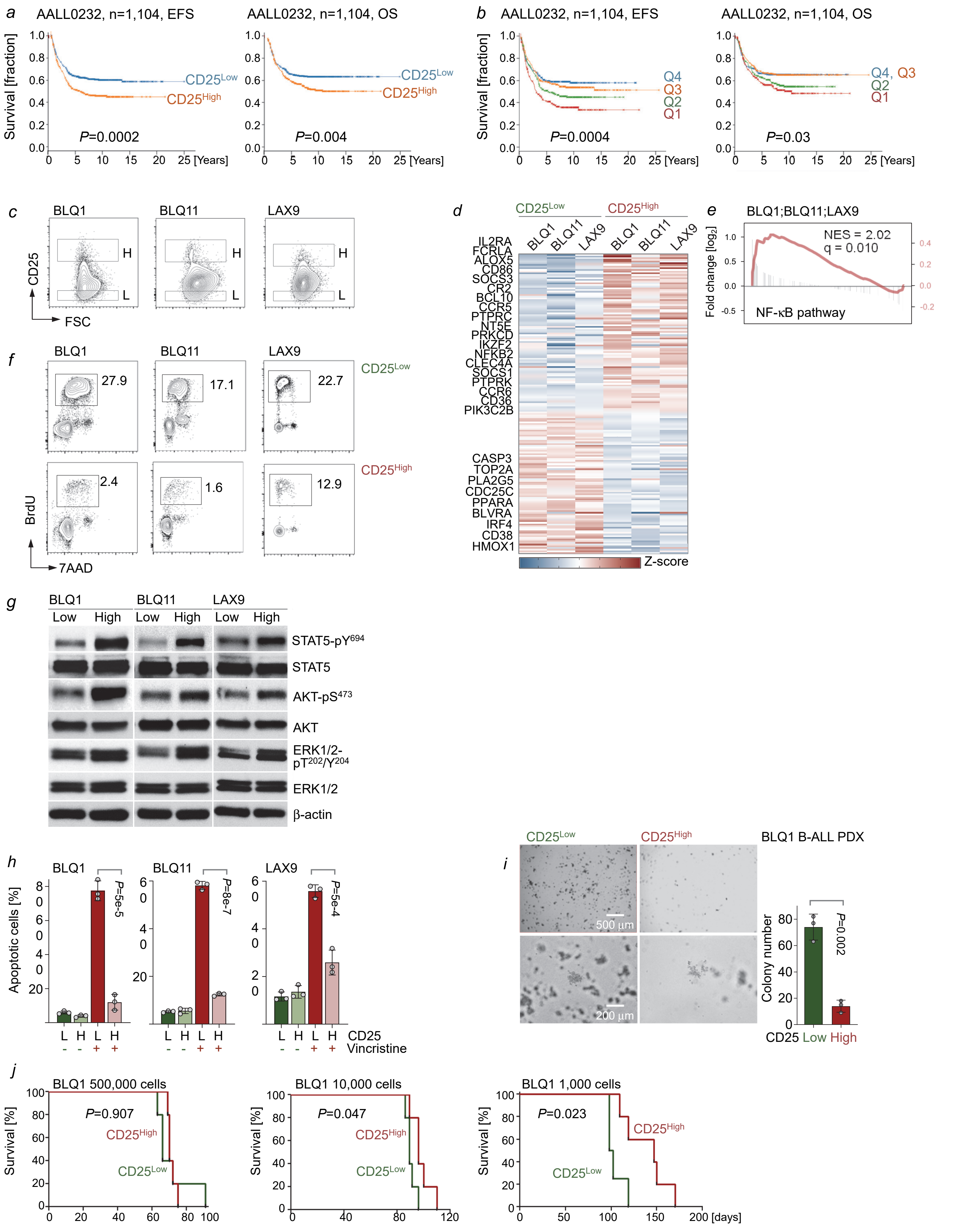

**Extended Data Figure 2:** *CD25 functions as a feedback regulator and promotes drug-resistance in patient-derived B-ALL*

Patients from the pediatric NCI-COG trial AALL0232 (n=1,104) were grouped based on higher than median or lower than median CD25 (IL2RA) mRNA levels (**a**) and event-free survival (EFS) and overall survival (OS) in the two groups (n=552 in each group) was compared by Kaplan-Meier analysis. In the same dataset, patients were assigned to four quartiles (Q1-Q4; n=276 in each quartile) based on CD25 mRNA levels and EFS and OS in the four groups was compared (**b**).

Levels of CD25 on surface were assessed by flow cytometry in each of three different patient-derived $Ph^+$ B-ALL cells (BLQ1, BLQ11, and LAX9). $CD25^{Low}$ (L) and $CD25^{High}$ (H) cells were sorted by flow cytometry (**c**) and gene expression was studied by Affymetrix GeneChips. The heatmap showed up- or down-regulated genes in $CD25^{High}$ versus $CD25^{Low}$ cells ($P<0.05$ and $\log_2$-transformed fold change >1; limma moderated *t*-statistic) (n=3; **d**).

Gene set enrichment analysis showed a significant correlation of gene signature of NF-κB pathway in $CD25^{High}$ versus $CD25^{Low}$ B-ALL cells. Red lines indicate running enrichment score (right-axis) and grey bars indicate log2 fold change (left-axis). Statistical significance was determined by two-tailed Kolmogorov-Smirnov test (**e**).

Cell cycle phases were analyzed by BrdU-incorporation in combination with 7-AAD staining in three patient-derived B-ALL cells, and the percentages of cells in S phase are indicated (**f**). Flow sorted $CD25^{High}$ or $CD25^{Low}$ cells from patient-derived B-ALL cells were studied by Western blots to measure STAT5-$pY^{694}$, STAT5, AKT-$pS^{473}$, AKT, ERK1/2-$pT^{202}/Y^{204}$ and ERK1/2 using β-actin as loading control (**g**).

Flow sorted $CD25^{High}$ or $CD25^{Low}$ cells from patient-derived B-ALL cells were treated with Vincristine (10 nM) or vehicle control for five days. The percentages of Annexin $V^+$ apoptotic cells quantitated by flow cytometry were plotted (**h**). Flow sorted $CD25^{High}$ or $CD25^{Low}$ cells from patient-derived $Ph^+$ B-ALL cells (SFO2) were subjected to colony forming assay. Total colony numbers were counted 14 days after plating in semi-solid methylcellulose. Representative microscopic images of colony formation assay are shown with colony numbers. Scale bars, 500 µm (top row), 200 µm (bottom row) (**i**).

Flow sorted $CD25^{High}$ or $CD25^{Low}$ cells from patient-derived B-ALL cells (BLQ1) were injected into NSG recipient mice with indicated cell numbers. Kaplan-Meier analyses were performed to compare overall survival of recipient mice (n=5 per group). Statistical significance was assessed by Mantel–Cox log-rank test (**j**).

In **f, g,** Statistical significance was assessed by two-tailed *t*-test (means ± s.d.). In **d-g,** Data are representative of three independent experiments.

**Extended Data Figure 3:** *Divergent outcome of CD25-deletion during early and late B-cell development*

*a*

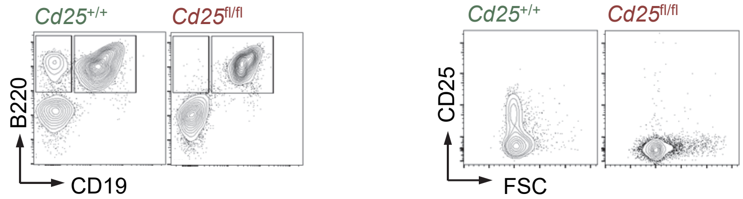


**a-b,** Expression of B220, CD19 and CD25 was analyzed by flow cytometry in bone marrow of *Cd25*$^{+/+}$ *Mb1*-Cre (n=7) and *Cd25*$^{fl/fl}$ *Mb1*-Cre (n=6) littermates (**a**). Proportion of early progenitors (B220$^{+}$, CD43$^{+}$), mature B-cells (B220$^{+}$, CD43$^{-}$), Hardy fraction C (B220$^{+}$, CD43$^{+}$, CD24$^{low}$, BP-1$^{high}$), fraction C' (B220$^{+}$, CD43$^{+}$, CD24$^{high}$, BP-1$^{high}$), fraction D (B220$^{+}$, CD43$^{-}$, IgM$^{low}$, IgD$^{low}$) and fraction F (B220$^{+}$, CD43$^{-}$, IgM$^{high}$, IgD$^{high}$) are shown **(b).**

*b*

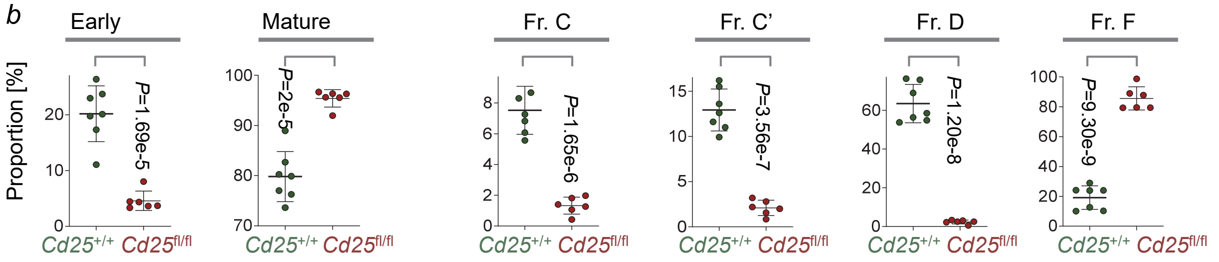


*c*

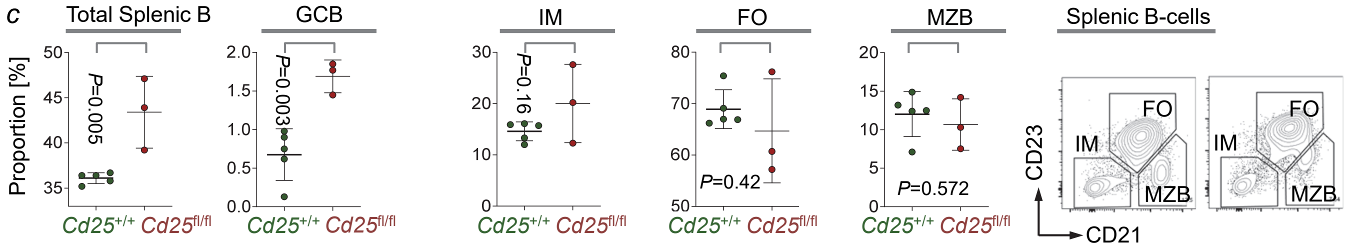


*d*

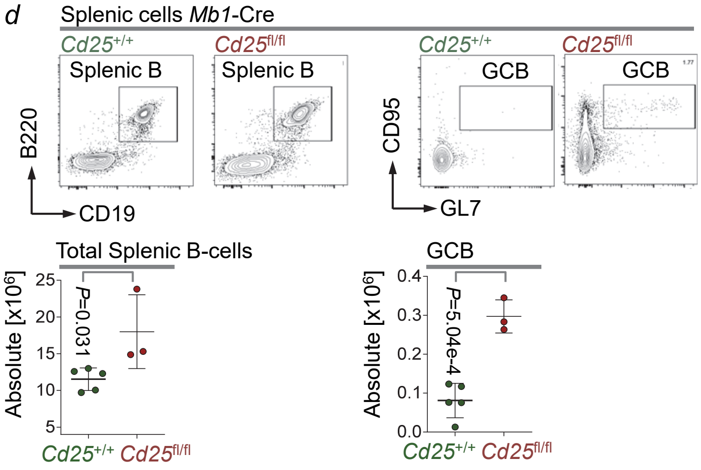


**c-e,** Proportion of total splenic B- (B220$^{+}$, CD19$^{+}$), germinal center B- (B220$^{+}$, CD19$^{+}$, CD95$^{+}$, GL7$^{+}$), immature B- (B220$^{+}$, CD19$^{+}$, CD23$^{low}$, CD21$^{low}$), follicular B- (B220$^{+}$, CD19$^{+}$, CD23$^{high}$, CD21$^{mid}$) and marginal zone B- (B220$^{+}$, CD19$^{+}$, CD23$^{low}$, CD21$^{high}$) cells from *Cd25*$^{+/+}$ *Mb1*-Cre (n=5) and *Cd25*$^{fl/fl}$ *Mb1*-Cre (n=3) littermates are plotted with flow cytometric analysis (**c**). Representative flow cytometry plots (top) for total splenic B- (B220$^{+}$, CD19$^{+}$) and germinal center B- (B220$^{+}$, CD19$^{+}$, CD95$^{+}$, GL7$^{+}$) cells are shown with absolute cell numbers (bottom) (**d**).

Absolute cell counts and proportions of CD19$^{+}$ total B-, B1 (CD19$^{+}$ Mac1+, IgM+), B1a (CD19$^{+}$, CD5$^{+}$, IgM$^{+}$) and B2 (CD19$^{+}$, B220$^{+}$, CD43$^{-}$, IgM$^{low}$) cells in the peritoneal cavity (PC) from *Cd25*$^{+/+}$ *Mb1*-Cre (n=5) and *Cd25*$^{fl/fl}$ *Mb1*-Cre (n=3) littermates are plotted with flow cytometric analysis (bottom). Representative flow cytometry plots (top) for B1 and total B-cells are shown. In **b-e,** two-tailed *t*-test (means ± s.d.).

*e*

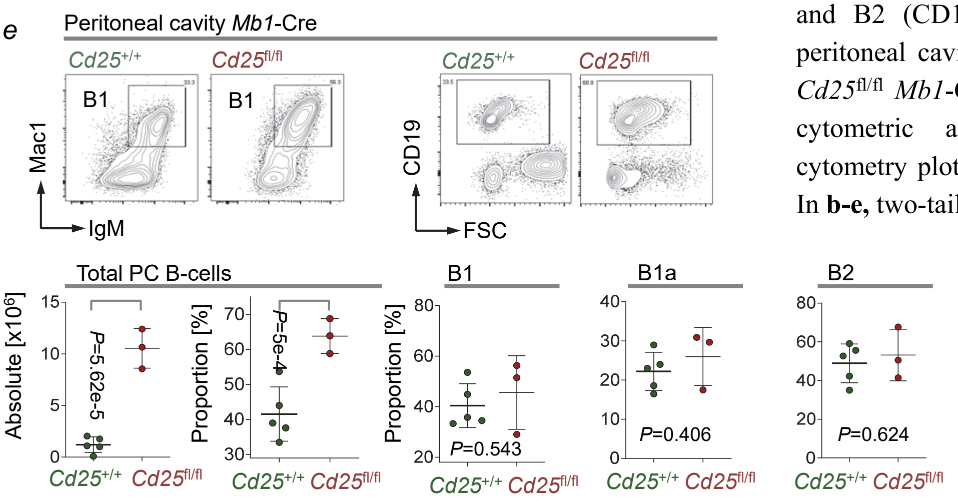

**Extended Data Figure 4:** *CD25 mediates negative feedback control of NF-κB signaling*

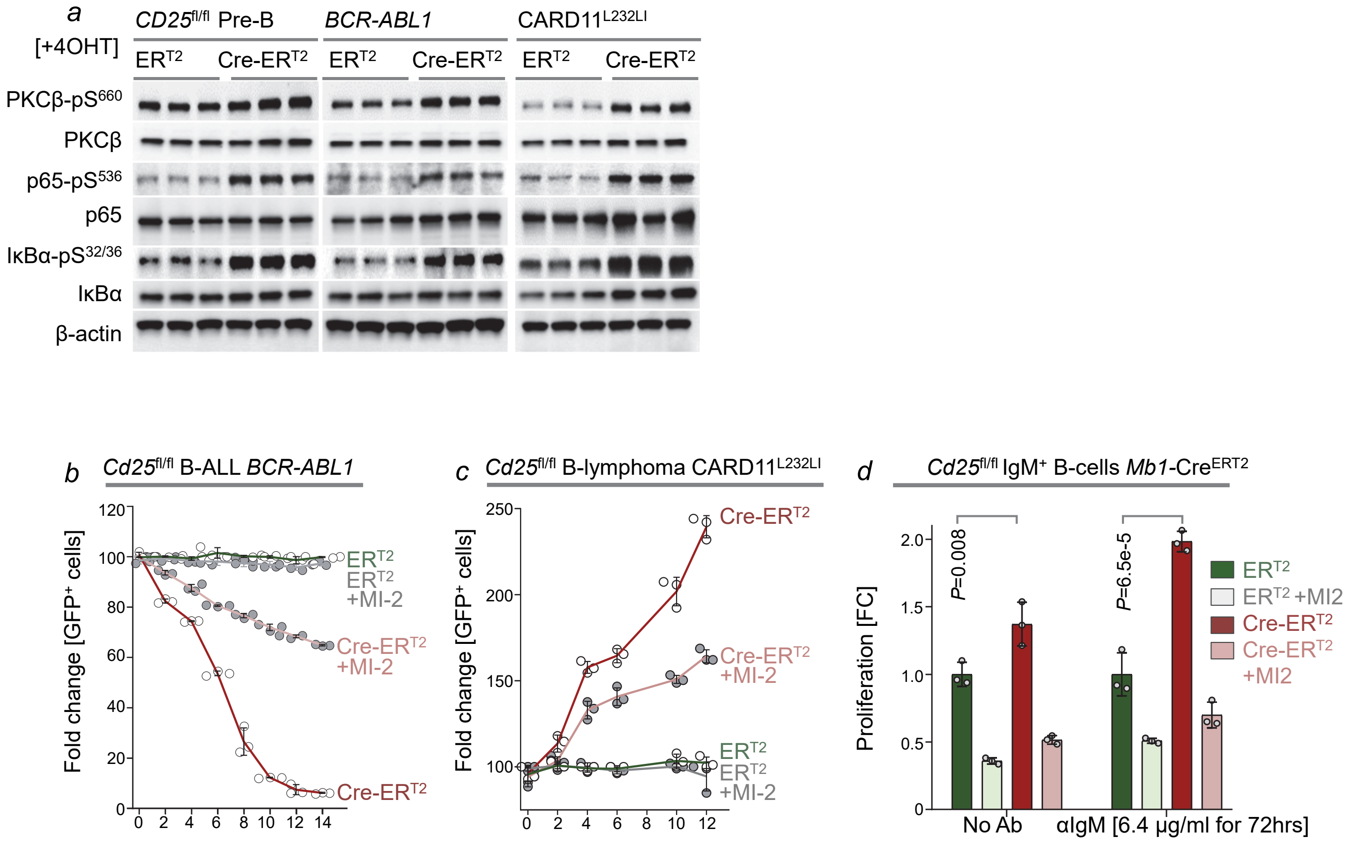


**a-b,** Expression of B220, CD19 and CD25 was analyzed by flow cytometry from bone marrow of *Cd25*$^{+/+}$ *Mb1*-Cre (n**a,** *Cd25*$^{fl/fl}$ IL7-dependent pre-B, *BCR-ABL1*-transformed B-ALL and *CARD11*$^{L232LI}$-transformed splenic mature B lymphoma cells carrying 4-OHT-inducible Cre-ER$^{T2}$ or ER$^{T2}$ were treated with 4-OHT for 2 days and studied by Western blot for protein levels of pan PKC (PKCβ)-pS$^{660}$, PKCβ, NF-κB p65-pS$^{536}$, NF-kB p65, IκBα-pS$^{32/36}$ and IκBα using β-actin as loading control.

**b,** *Cd25*$^{fl/fl}$ B-ALL cells transformed with BCR-ABL1 were transduced with 4-OHT-inducible Cre-ER$^{T2}$-GFP or ER$^{T2}$-GFP empty vector control. Percentages of GFP$^{+}$ cells were measured by flow cytometry at different time points following 4-OHT treatment in the presence of MI-2 (500 nM) or vehicle control.

**c,** *Cd25*$^{fl/fl}$ mature B cells purified from splenocytes by negative selection were retrovirally transformed with CARD11$^{L232LI}$-mCherry and then transduced with 4-OHT-inducible Cre-ER$^{T2}$-GFP or ER$^{T2}$-GFP in the presence of IL-4 (10 ng/ml), BAFF (100 ng/ml) and CD40L (1 mg/ml). Enrichments of GFP$^{+}$mCherry$^{+}$ B-cell lymphoma cells were measured by flow cytometry at different time points following 4-OHT treatment in presence of MI-2 (50 nM) or vehicle control.

**d,** Mature B cells negatively purified from splenocytes of *Cd25*$^{fl/fl}$ *Mb1*-Cre-ER$^{T2}$ mice were suspended in complete medium in the absence of Il-4, BAFF and CD40-ligand and treated with anti-IgM antibody or isotype control in combination with 4-OHT, MI-2 (500 nM) or vehicle control for 3 days and relative proliferation was assessed by CellTiter-Glo assay. In **a-d,** Data are representative of three independent experiments. In **d,** mean ± s.d. indicated; significance determined by two-tailed *t*-test.

**Extended Data Figure 5:** *CD25 functions as a feedback regulator of BCR-downstream signaling*

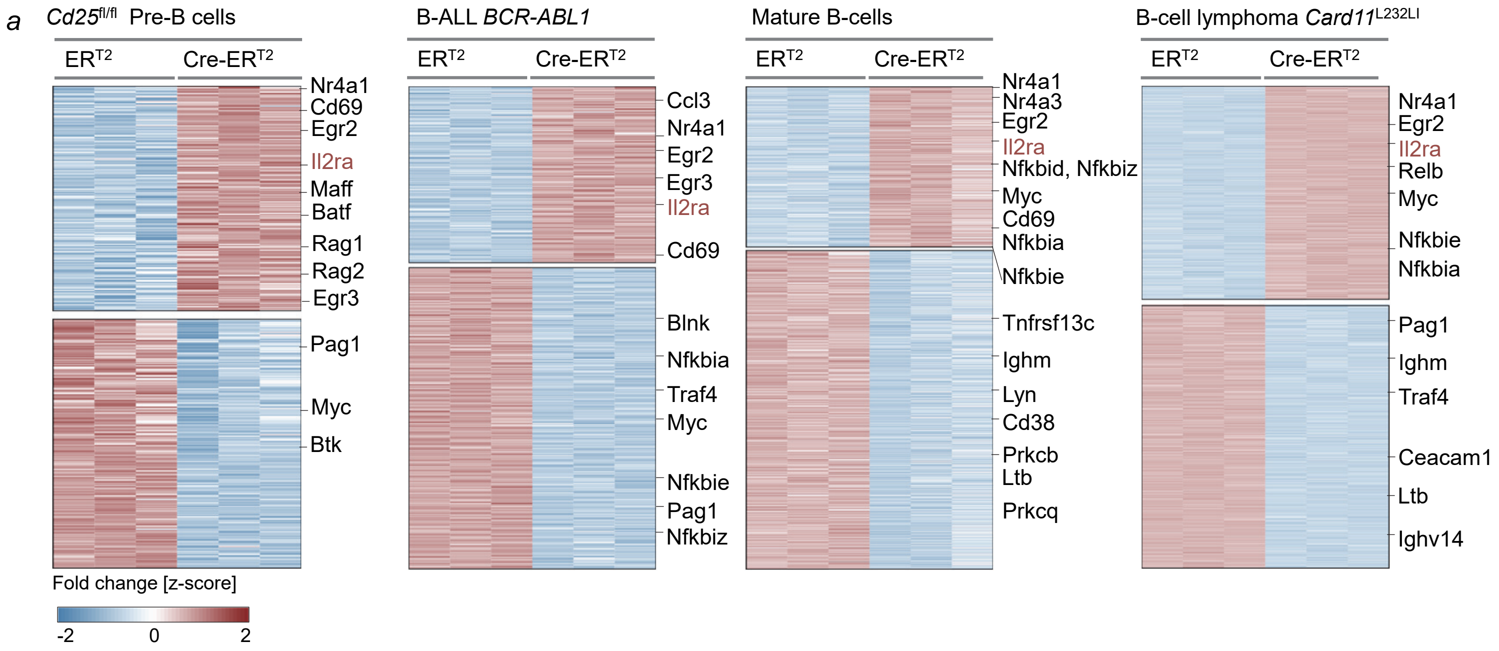


**a-c,** RNA sequencing was performed with *Cd25*fl/fl IL7-dependent pre-B, *BCR-ABL1*-transformed B-ALL, splenic mature B and *CARD11*L232LI-transformed splenic B lymphoma cells carrying 4-OHT-inducible Cre-ERT2 or ERT2 upon treatment of 4-OHT or vehicle for 2 days. Relative rlog-normalized gene expression values for all strongly differentially expressed genes ($P<0.05$ and $\log_2$-transformed fold change >1; Wald test with Benjamini–Hochberg correction) plotted as a heat map with row scaling. Upregulation of *Il2ra* was highlighted in red (**a**).

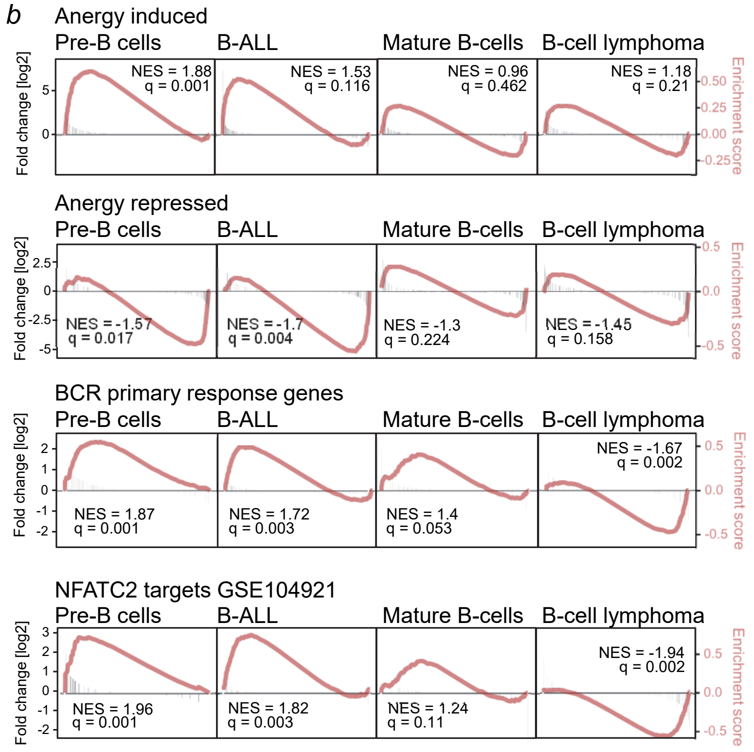


Gene set enrichment analysis for genes ranked by ratio of 4-OHT to vehicle as $\log_2$-transformed fold change. Red lines indicate running enrichment score (right axis), grey bars indicate log2 fold change (left axis). Statistical significance was determined by two-tailed Kolmogorov-Smirnov test (**b**).

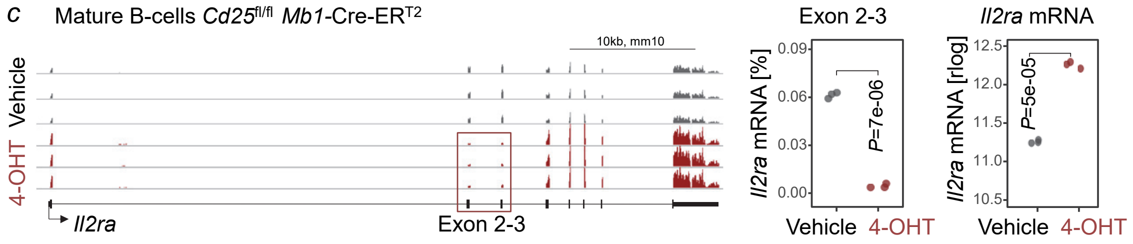


Reads aligned to the *Cd25* locus, floxed exons 2-3 are highlighted (left). The relative percentage of total *Cd25*-aligned reads mapped to floxed exons 2-3 (middle) and rlog normalized expression of total *Cd25* transcripts are plotted (right). Statistical significance was assessed by two-tailed *t*-test (**c**).

**Extended Data Figure 6:** *CD25 prevents formation of spontaneous germinal centers and expansion of autoreactive B-cells*

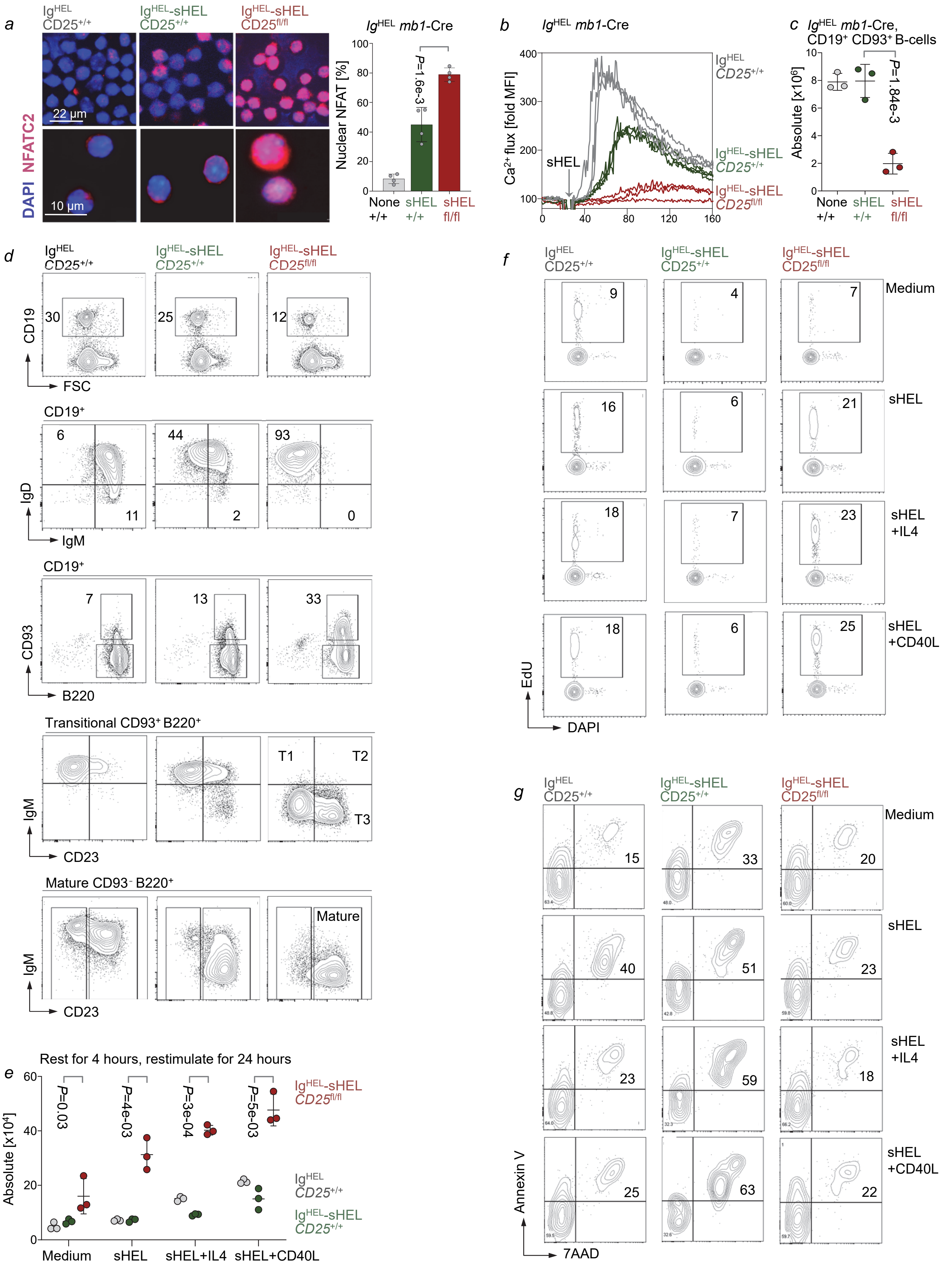

**Extended Data Figure 6:** *CD25 prevents formation of spontaneous germinal centers and expansion of autoreactive B-cells*

**a-b,** Mature B-cells from splenocytes of *Cd25*$^{+/+}$ Ig$^{HEL}$, *Cd25*$^{+/+}$ Ig$^{HEL}$-sHEL *Mb1*-Cre and *Cd25*$^{fl/fl}$ Ig$^{HEL}$-sHEL *Mb1*-Cre mice were purified by negative selection and NFATC2 localization was analyzed by immunofluorescence with NFATC2 and DAPI staining (left). Bar graph (right) shows percentages of nuclear NFATC2 with mean ± standard error analyzed using QuPath (**a**).

$Ca^{2+}$ mobilization in response to 0.25 μg/ml soluble HEL (sHEL) engagement was measured using cell-permeant Flou-4 dye (**b**). **c-d,** Mature B-cell subsets isolated from spleen of *Cd25*$^{+/+}$ IgHEL (n=3), *Cd25*$^{+/+}$ IgHEL-sHEL *Mb1*-Cre (n=3) or *Cd25*$^{fl/fl}$ IgHEL-sHEL *Mb1*-Cre (n=3) mice were analyzed by flow cytometry and absolute cell counts of transitional B-cells are plotted (**c**). CD19, IgD, CD23 and IgM expression was assessed by flow cytometry to identify T1 (CD19$^{+}$, B220$^{+}$, CD93$^{+}$, IgM$^{high}$, CD23$^{-}$), T2 (CD19$^{+}$, B220$^{+}$, CD93$^{+}$, IgM$^{high}$, CD23$^{+}$), T3 (CD19$^{+}$, B220$^{+}$, CD93$^{+}$, IgM$^{low}$, CD23$^{+}$) and mature (CD19$^{+}$, B220$^{+}$, CD93$^{-}$, CD23$^{+}$) populations (**d**).

**e-g,** Mature B-cells purified from splenocytes by negative selection were incubated for 4 hours in the absence of soluble HEL antigen and restimulated with soluble HEL antigen (sHEL, 0.5 μg/ml) in combination with IL4 (10 ng/ml) or CD40 ligand (CD40L, 10 ng/ml) for 24 hours and the number of viable cells were counted (**e**). Cell cycle analyses were performed by measuring EdU incorporation in combination with DAPI staining, and the percentages of cells in S phase are indicated (**f**). Cell viability was measured by Annexin V/7AAD staining, and the percentages of cells with Annexin V$^{+}$ 7AAD$^{+}$ are indicated (**g**).

In **a,b, e-g,** Data are representative of three independent experiments. In **c, e,** mean ± s.d. indicated; significance determined by two-tailed *t*-test.

**Extended Data Figure 7:** *Developmental defects of CD25-deficient B-cells are not related to defective IL2 signaling*

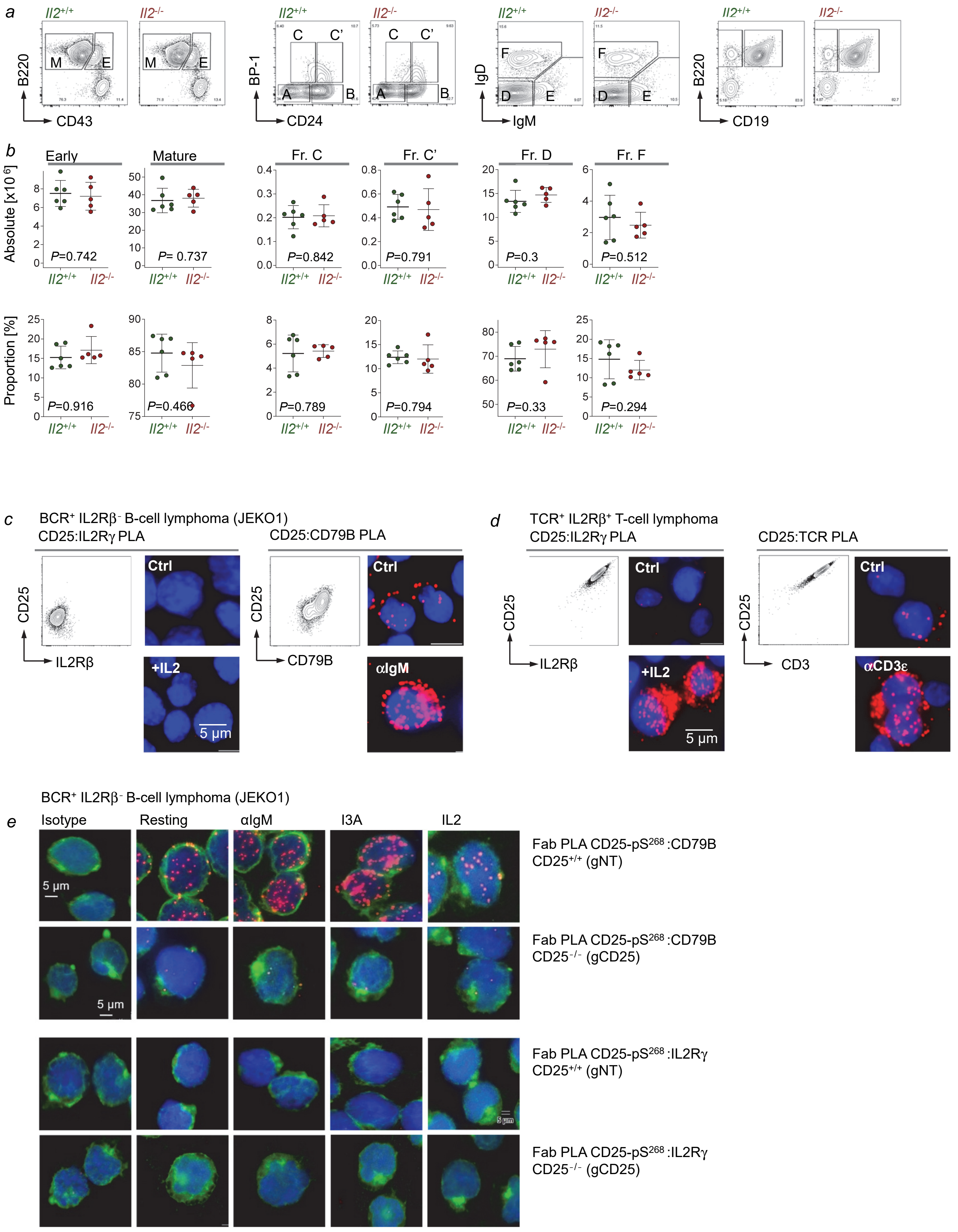

**Extended Data Figure 7:** *Developmental defects of CD25-deficient B-cells are not related to defective IL2 signaling*

**a-b,** Hardy fractions of B cell subsets isolated from bone marrow of *Il2*$^{+/+}$ (n=6) and *Il2*$^{-/-}$ (n=5) littermates were analyzed by flow cytometry. In Nk1.1$^{-}$ and Gr1$^{-}$ populations, absolute cell counts and proportion of early progenitor (B220$^{+}$, CD43$^{+}$), mature (B220$^{+}$, CD43$^{-}$), fraction C (B220$^{+}$, CD43$^{+}$, CD24$^{low}$, BP-1$^{high}$), fraction C' (B220$^{+}$, CD43$^{+}$, CD24$^{high}$, BP-1$^{high}$), fraction D (B220$^{+}$, CD43$^{-}$, IgM$^{low}$, IgD$^{low}$) and fraction F (B220$^{+}$, CD43$^{-}$, IgM$^{high}$, IgD$^{high}$) cells are plotted (means ± s.d.; two-tailed t-test). Representative flow cytometry plots are shown.

**c-d,** Surface expression of CD25, IL2Rβ and CD79B in JEKO1 cells was measured by flow cytometry. The proximity of CD25 to IL2Rγ or CD79B was assessed by *in situ* Fab Proximity ligation assay (PLA) as previously described[1] in JEKO1 cells stimulated with IL2 (10 ng/ml) or anti-IgM (F(ab')$_2$ fragments of anti-human μ chain, 10 μg/ml) for 5 min, respectively (**c**). Surface expression of CD25, IL2Rβ and CD3ε in Fe-PD T-cell lymphoma cells was measured by flow cytometry. The proximity of CD25 to IL2Rγ or CD3δ was assessed after stimulation of Fe-PD cells with IL2 (10 ng/ml) or anti-CD3ε (10 μg/ml) for 5 min (**d**).

**e,** The proximity of CD25-pS$^{268}$ to CD79B or IL2Rγ was assessed by *in situ* Fab PLA as previously described[1] in non-targeted JEKO1 (gNT) cells stimulated with anti-IgM (F(ab')$_2$ fragments of anti-human μ chain, 10 μg/ml), I3A (50 nM) or IL2 (10 ng/ml) for 5 min. The isotype control for CD79B antibody and *CD25*$^{-/-}$ JEKO1 (gCD25) cells with CRISPR-Cas9-mediated gene deletion was used as negative control.

In **c-e**, PLA signal is shown as red dot and nuclei stained with DAPI as blue and representative microscopic images are shown (n=3). Scale bar, 5 μm.

**Reference**

[1]Kläsener K, Yang J, Reth M. Study B Cell Antigen Receptor Nano-Scale Organization by In Situ Fab Proximity Ligation Assay. *Methods Mol Biol.* 2018; 1707:171-181.

**Extended Data Figure 8:** *Lack of IL2Rβ chain expression explains IL2-unresponsiveness in B-cells*

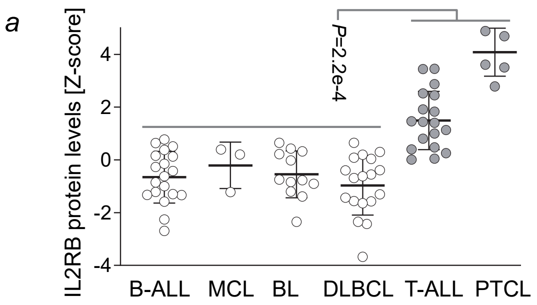


**a,** Protein levels of IL2Rβ were assessed in different types of B-ALL, MCL, BL, DLBCL and compared with T-ALL and PTCL.

**b,** Eleven different B-ALL, MCL, DLBCL, Multiple Myeloma and PTCL cell lines (**Supplementary Table 3**) were analyzed by Western blot for levels of STAT5-pY$^{694}$, STAT5, ERK1/2-pT$^{202}$/Y$^{204}$ and ERK1/2 upon stimulation with IL2 (10 ng/ml) or vehicle control for 5 min. β-actin was used as loading control.

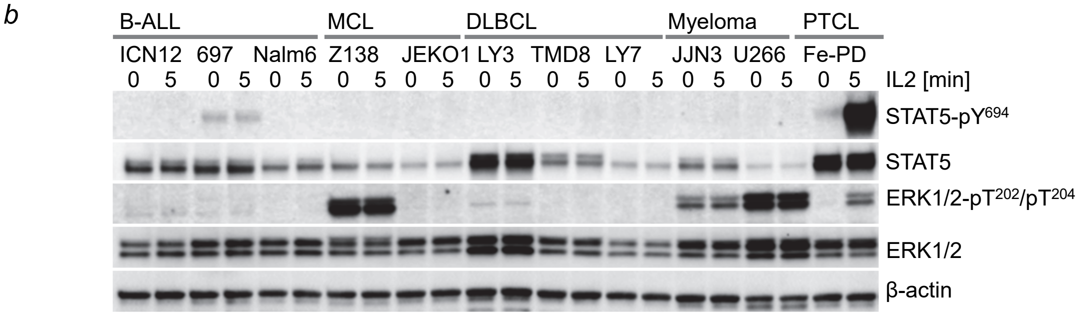


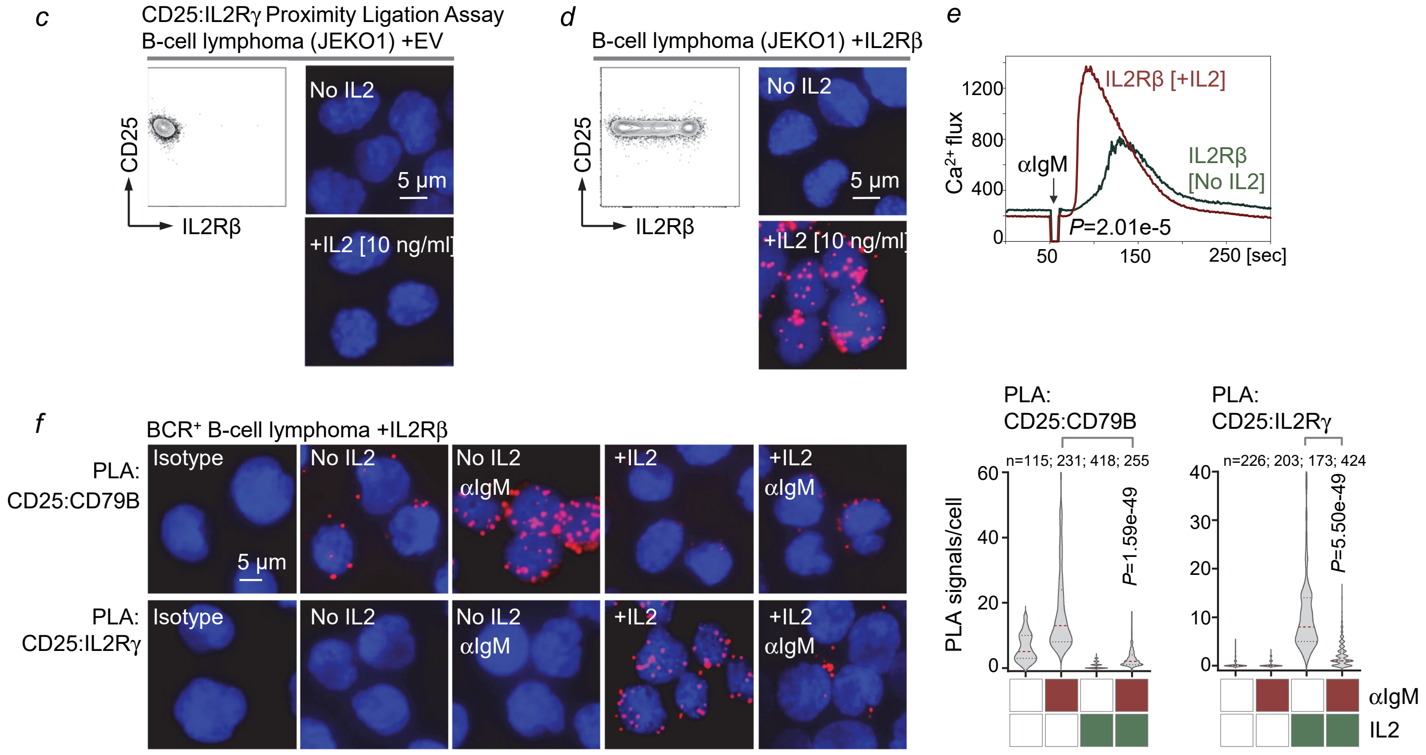


**c-f**, Surface expression of CD25 and IL2Rβ in JEKO1 cells transduced with empty vector control (EV) (**c**) or IL2Rβ (**d**) was measured by flow cytometry. The proximity of CD25 to IL2Rγ was assessed upon stimulation with IL2 (10 ng/ml) or vehicle control for 5 min.

**e**, JEKO1 cells transduced with IL2Rβ were incubated with IL2 (10 ng/ml) or vehicle control for 5 min prior to Flou-4 loading. $Ca^{2+}$ mobilization in response to anti-IgM (F(ab')$_2$ fragments of anti-human μ chain, 10 μg/ml) mediated BCR engagement was measured by flow cytometry.

**f,** JEKO1 cells transduced with IL2Rβ were stimulated for 5 min with anti-IgM (F(ab')$_2$ fragments of anti-human μ chain, 10 μg/ml) in the presence or absence of IL2 (10 ng/ml). The proximity of CD25:CD79B (top) and CD25:IL2Rγ (bottom) was examined and shown as representative microscopic images (left) and quantified (right).

In **c, d, f,** Scale bars, 5 μm. In **a, e, f,** mean ± s.d. indicated; significance determined by two-tailed *t*-test. **a-f,** Data are representative from three independent experiments. For gel source data, see **Supplementary Fig.1**.

**Extended Data Figure 9:** *Recruitment of CD25 reflects different types of oncogenic BCR-signaling*

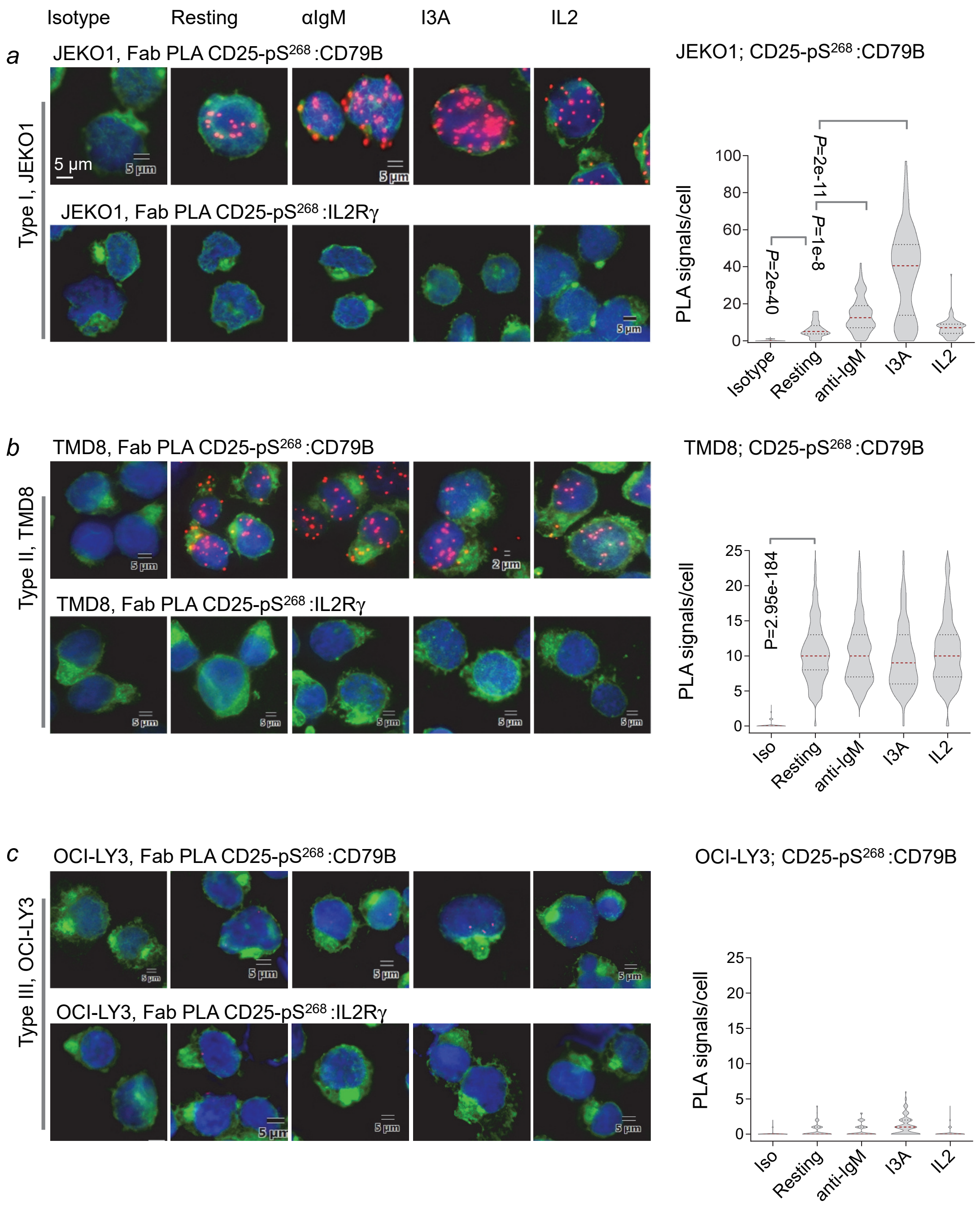


**a-c,** The proximity of CD25-pS268 to CD79B or IL2Rγ was assessed by *in situ* Fab PLA as previously described[1] in each of three different types of lymphoma cells, characterized by BCR signalling as JEKO1 MCL (inducible BCR), TMD8 ABC-DLBCL (chronic active BCR) and OCI-LY3 ABC-DLBCL (BCR-independent). Cells were stimulated with anti-IgM (F(ab')$_2$ fragments of anti-human µ chain, 10 µg/ml), I3A (50 nM) or IL2 (10 ng/ml) for 5 min. The isotype control for CD79B antibody was used as negative control (left). Quantitative analysis (right) shown as Fab PLA signals per cell of each sample with the mean values as red line. Significance determined by two-tailed *t*-test. Scale bars, 5 µm. Data are representative from three independent experiments.

**Reference**


[1]Kläsener K, Yang J, Reth M. Study B Cell Antigen Receptor Nano-Scale Organization by In Situ Fab Proximity Ligation Assay. *Methods Mol Biol.* 2018; 1707:171-181.

**Extended Data Figure 10:** *Inherited PRKCD mutations in two families*

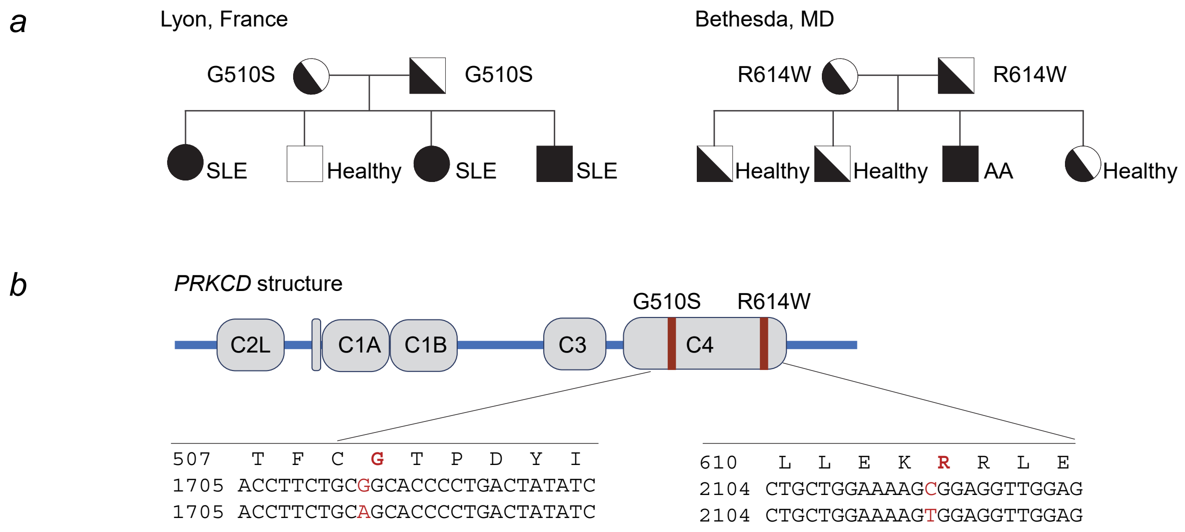


(**a**) Family pedigrees for two families (Bethesda and Lyon) with inherited *PRKCD* mutations (R614W and G510S) are shown. Siblings with homozygous mutations had autoimmune disease (AA, Autoantibody; SLE, systemic lupus erythematosus). (**b**) Schematic diagram of PKCδ domain structure annotated with location of the two mutations.

**Extended Data Figure 11:** *Structural modeling of RACK1:CD25:PKCδ complexes*

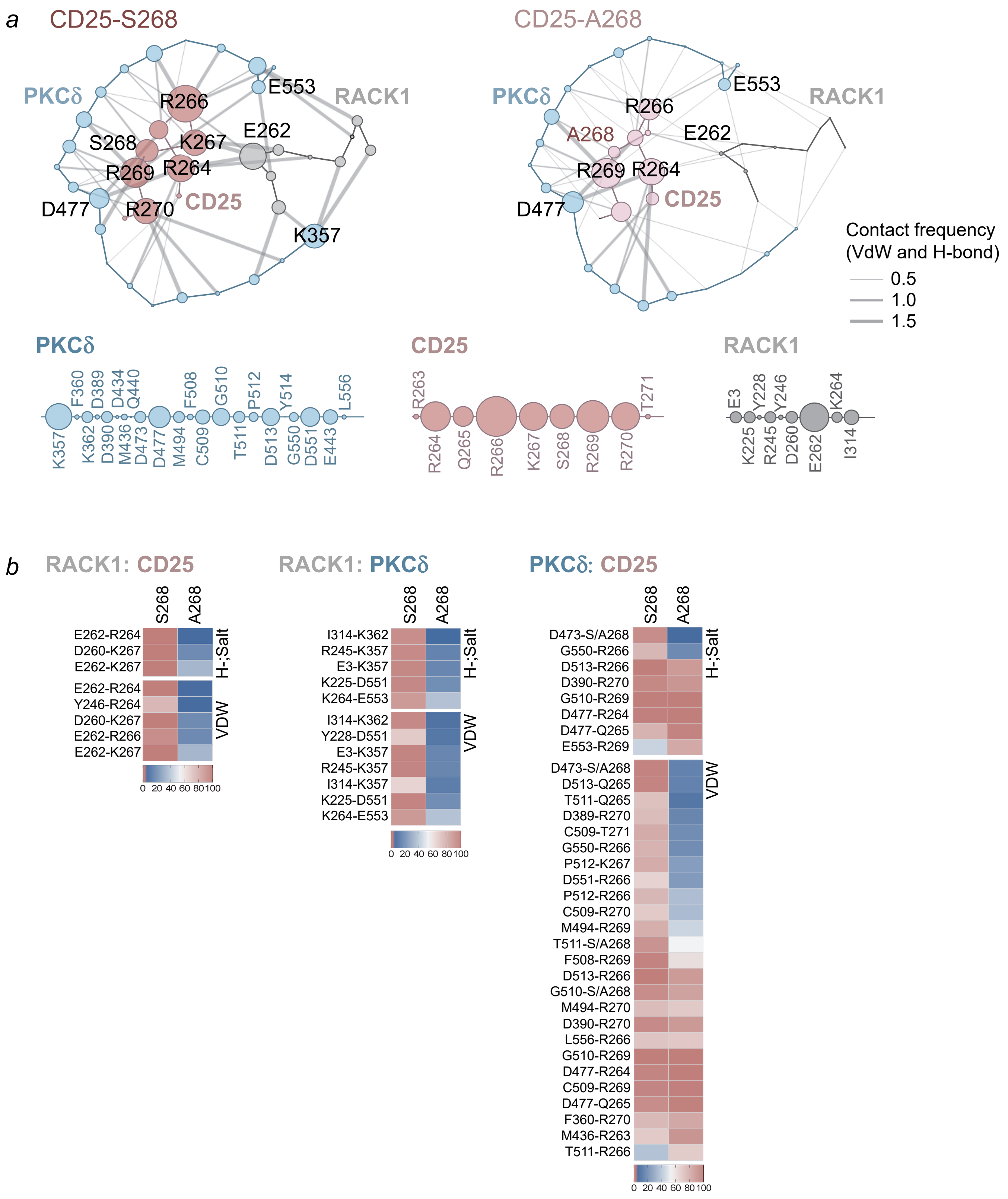


**a-b,** Multilayer residue-residue contact network shows multiple interactions between indicated amino acid residues of PKCδ, RACK1 and CD25 via van der Waals force (VDW), Hydrogen bond (H-bond) and salt-bridge. Size of circles and link thickness are proportional to the total number of interactions quantitated by the accumulated contact frequency in ternary RACK1:CD25:PKCδ complex (**a**). Heatmap of residue contact frequency was quantitated as the percentage of MD simulation frames that show the interaction in ternary RACK1:CD25:PKCδ complex and its mutant S268A. The total number of MD frames for each system is 25,000 (**b**).

**Extended Data Figure 12:** *CD25 functions as a feedback regulator of oncogenic RTK-signaling in myeloid leukemia cells*

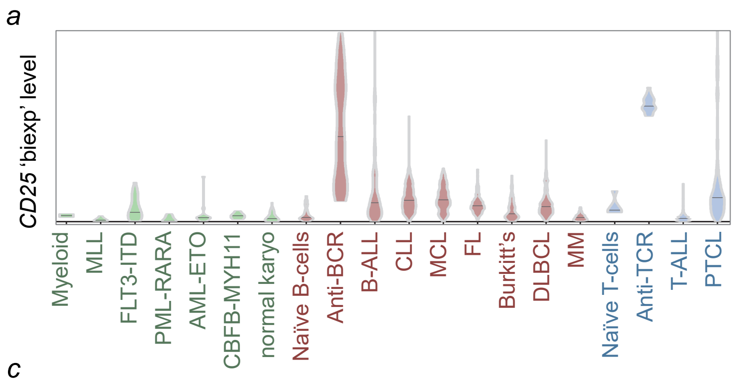


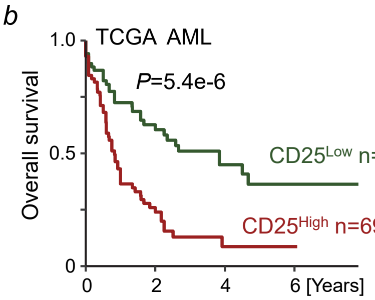


**a,** *CD25* mRNA levels in human normal and malignant myeloid, B- and T-lymphoid samples are plotted (**Supplementary Table 9**). **b,** Patients with AML (n=140, TCGA) were divided into two groups based on higher or lower than median levels of *CD25* mRNA. Overall survival was estimated in the two groups by Kaplan-Meier survival analyses. Log-rank test was used to assess statistical significance.

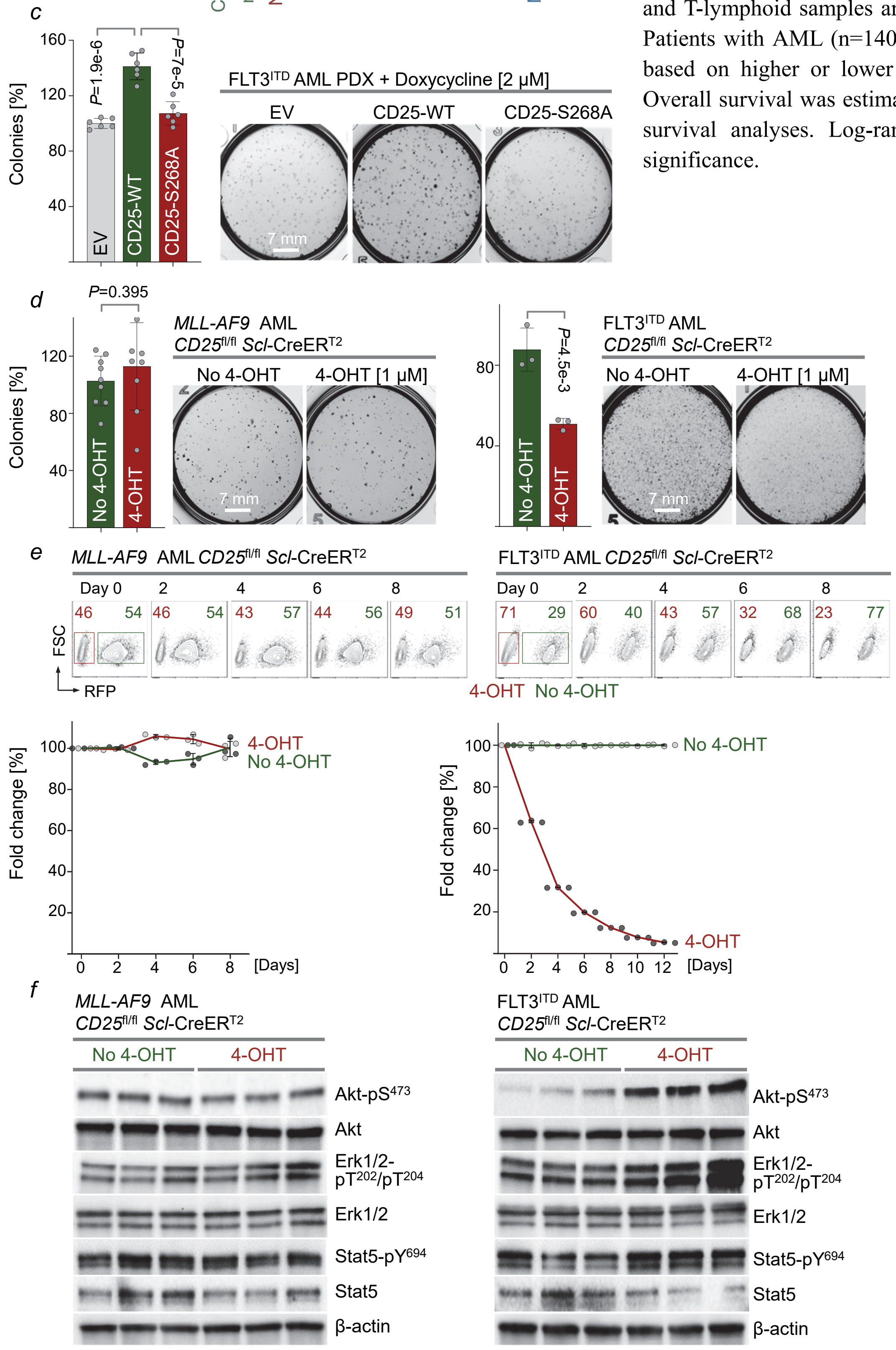


**c,** MLL-AF9$^+$ human CD34$^+$ cells were transformed with FLT3$^{ITD}$-GFP and GFP$^+$ cells were sorted by flow cytometry and transduced with doxycycline-inducible wild-type (CD25-WT), S268A mutant or empty vector (EV) and plated in semi-solid methylcellulose for colony-forming assay in the presence of 2 μM doxycycline. Total colony numbers were counted on day 8 after plating and relative colonies to EV control were plotted.

Myeloid progenitors from bone marrow of *Cd25*$^{fl/fl}$ *MLL-AF9*$^{Pos/Wt}$ *Scl*-Cre-ER$^{T2}$ mice were transformed with FLT3$^{ITD}$-GFP. Flow sorted GFP$^-$ (left) and GFP$^+$ (right) cells were plated in semi-solid methylcellulose for colony-forming assay in the presence of 2 μM 4-OHT or vehicle control as indicated. Relative colony numbers to EV control were assessed on day 7 (**d-e**).

Flow sorted GFP$^-$ (left) and GFP$^+$ (right) cells from (**d**) were treated with 4-OHT or vehicle control for 2 days and levels of Akt-pS$^{473}$, Akt, Erk1/2-pT$^{202}$/Y$^{204}$, Erk1/2, Stat5-pY$^{694}$ and Stat5 were assessed by Western blot with β-actin as loading control. In **c, d,** Significance determined by two-tailed *t*-test. (means ± s.d.), scale bars, 7 mm. In **c-f,** Data are representative from three independent experiments.

**Extended Data Figure 13:** *Differential outcome of CD25-targeting in B-ALL and mature B-cell lymphoma*

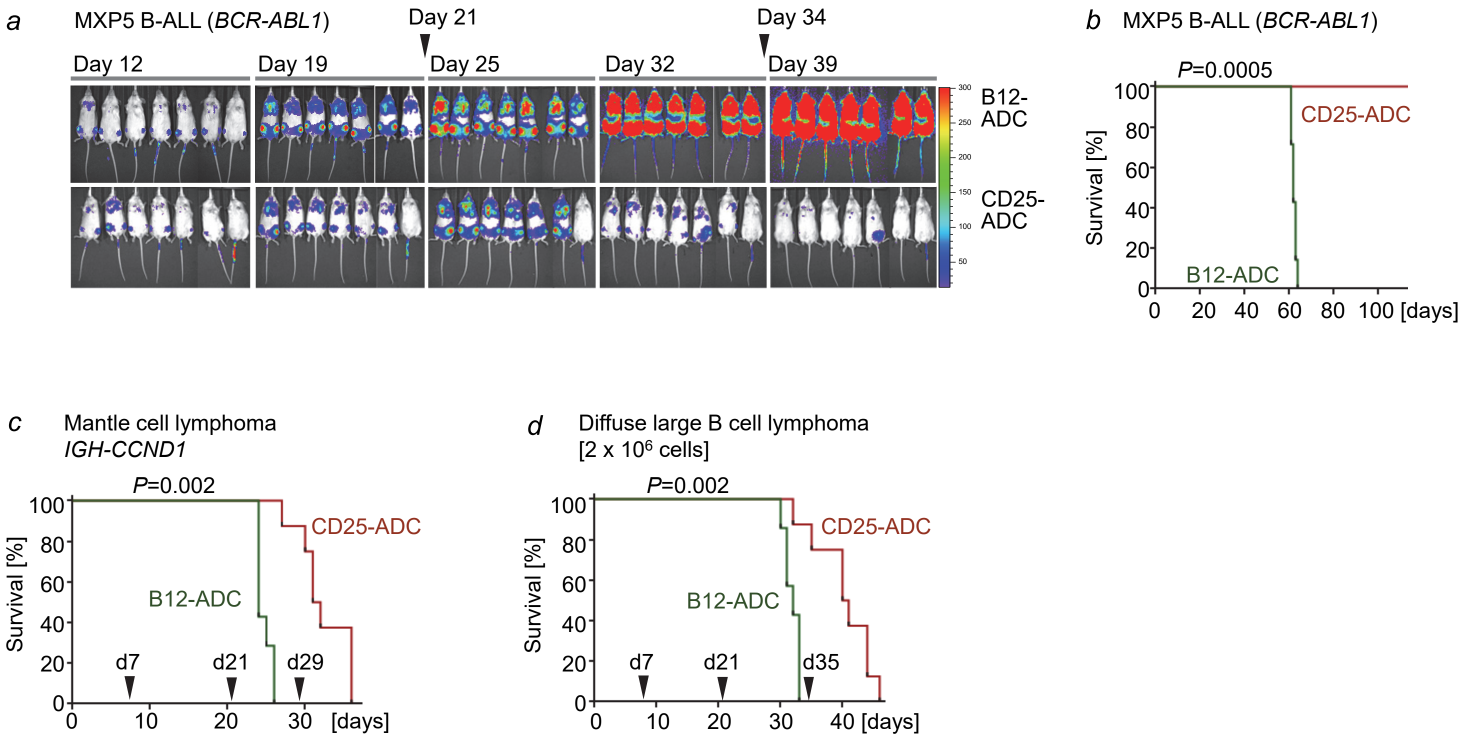


**a-d**, Patient-derived B-ALL (MXP5) cells were labeled with firefly luciferase and $2 \times 10^6$ cells were injected into sublethally irradiated (200 cGy) NSG recipient mice (n=7 per group). CD25-ADC, a CD25-targeted pyrrolobenzodiazepine (PBD) dimer-based antibody-drug conjugate (ADC), or the control ADC (B12-ADC) dissolved in PBS was administered intravenously (i.v.) at dose of 600 µg $kg^{-1}$ body weight at various time points as indicated by arrowheads.

The *in vivo* expansion and leukemia burden was measured by luciferase bioimaging at the indicated time points (**a**) and overall survival of recipient mice was determined by Kaplan-Meier analysis (**b**). Kaplan-Meier analyses of recipient mice (n=7, B12-ADC; n=8, CD25-ADC) injected with $2 \times 10^6$ cells of MCL PDX harboring *IGH-CCND1* (**c**) or DLBCL OCI-LY7 (**d**). ADCs (0.6 mg/kg) were administered intravenously (i.v.) at various time points as indicated by arrowheads. **b-d,** The *p* value was calculated with Gehan-Breslow-Wilcoxon test.
